# The KMT2A-MENIN complex is a therapeutic vulnerability in SETBP1-mutant leukemia

**DOI:** 10.64898/2026.11.17.765711

**Authors:** Samantha Tauchmann, Thai T Nguyen, Hanqian L Carlson, Amara Pang, Theodore P Braun, Julia E Maxson

## Abstract

SET binding protein 1 (*SETBP1*) is recurrently mutated in high-risk myeloid malignancies, including myelodysplastic/myeloproliferative overlap disorders (MDS/MPNs) and is associated with poor clinical outcomes. Although *SETBP1* mutations promote self-renewal and HOX-driven transcription in murine and cell line models, the epigenetic dependencies sustaining these programs in human hematopoietic stem and progenitor cells (HSPCs) remain poorly defined. Here, we identify the KMT2A-MENIN complex as a targetable dependency in *SETBP1*-mutant leukemia. Expression of *SETBP1*^*D868N*^ enhanced growth and serial replating of human HSPCs and established a transcriptionally permissive chromatin state, with increased H3K4me3, H3K27ac and KMT2A occupancy at *SETBP1*-regulated loci. Integrated transcriptional and chromatin profiling defined a 94-gene SETBP1 regulome containing *HOXA9, MEIS1, MECOM*, and *MYB* and was enriched for transcriptional programs associated with *KMT2A-*rearranged, *NUP98-HOXA9*, and *NPM1*-mutant leukemias. CRISPR-Cas9 disruption revealed a functional requirement for KMT2A, but not for KMT2B or DOT1L in *SETBP1*^*D868N*^ HSPCs. Strikingly, deletion of the KMT2A cofactor MEN1 phenocopied KMT2A loss. Disruption of either KMT2A or MEN1 suppressed *SETBP1*-driven proliferation and leukemogenic transcriptional programs, including leukemia stem cell-associated signatures. We next exploited this dependency pharmacologically. A KMT2A-MENIN inhibitor suppressed canonical *SETBP1* target genes and abolished mutant *SETBP1*-driven serial replating while promoting hematopoietic differentiation and reducing H3K4me3 at leukemogenic loci, despite modest effects on short-term proliferation. Most importantly, three clinically relevant KMT2A-MENIN interaction inhibitors impaired clonogenic growth across primary *SETBP1*-mutant leukemia patient samples, while inducing myeloid differentiation and suppressing *SETBP1* target gene expression. Together, these findings identify the KMT2A-MENIN complex as a key therapeutic vulnerability in *SETBP1*-mutant leukemia. The consistent activity of multiple MENIN inhibitors across a variety of *SETBP1*-mutant patient samples and cell models validates the KMT2A-MENIN interaction as an actionable vulnerability and provides a strong rationale for expanding MENIN-directed therapies to *SETBP1*-mutant myeloid malignancies.

**KEYPOINTS:**

- Mutant *SETBP1* activates a KMT2A-MENIN-dependent leukemogenic transcriptional and chromatin program
- KMT2A-MENIN inhibition suppresses self-renewal and induces differentiation, exposing a therapeutic vulnerability in *SETBP1*-mutant leukemia

## INTRODUCTION

SET binding protein 1 (*SETBP1*) is recurrently mutated in several high-risk myeloid malignancies and is particularly enriched in myelodysplastic/myeloproliferative overlap disorders (MDS/MPNs). *SETBP1* mutations are particularly common in chronic neutrophilic leukemia (CNL, ∼50%), and have been reported across multiple MDS/MPN entities, including MDS/MPN with neutrophilia (MDS/MPN-N, formerly atypical CML, ∼32%), chronic myelomonocytic leukemia (CMML, ∼18%), juvenile myelomonocytic leukemia (JMML, ∼24%), and unclassifiable MDS/MPN (MDS-MPN-U, ∼10%) cases^1–8^. These mutations cluster within a hotspot region that disrupts protein degradation, leading to increased SETBP1 protein abundance^4^. Across disease subtypes, *SETBP1* mutations are consistently associated with reduced overall survival, increased risk of relapse, and therapy resistance^1,7,9–11^. Despite their strong prognostic significance, no targeted therapies currently exist for *SETBP1-*mutant leukemias.

Prior studies, largely conducted in murine progenitor systems, demonstrated that Setbp1 overexpression promotes self-renewal and activate HOX-associated transcriptional programs, including *Hoxa9* and *Hoxa10*^12–14^. Additional murine models have shown that mutant *SETBP1* cooperates with other oncogenic lesions to accelerate aggressive myeloid disease and reinforce aberrant transcriptional and epigenetic states^15,16^. While these studies establish the oncogenic potential of increased *SETBP1* expression, the chromatin-regulatory mechanisms required to sustain human *SETBP1*-driven transcriptional programs remain incompletely defined.

Recent work suggests that *SETBP1*-driven leukemias may converge on epigenetic dependencies shared with other high-risk myeloid malignancies. *SETBP1*-mutant cells exhibit transcriptional signatures enriched for canonical stemness regulators, including *HOXA* cluster genes, *MEIS1*, and *MYC-*associated programs^12,15,17,18^. Strikingly, KMT2A-associated transcriptional programs converge on activation of *HOXA/MEIS1* gene networks that are essential for leukemic self-renewal^19,20^. Moreover, KMT2A physically interacts with both wild-type and mutant SETBP1, establishing the KMT2A complex as an essential regulator of gene expression and the transformation of hematopoietic stem and progenitor cells (HSPCs)^17,18,21^. In *KMT2A*-rearranged and *NPM1*-mutant leukemias, leukemogenic transcriptional programs depend on the interaction between KMT2A and MENIN, a chromatin-associated scaffolding protein encoded by *MEN1* that stabilizes KMT2A binding at target loci^22^. Pharmacologic disruption of this interaction collapses leukemogenic gene expression, induces differentiation, and has demonstrated clinical efficacy^23–26^. However, whether *SETBP1*-mutant leukemias similarly depend on the KMT2A-MENIN complex remains unknown.

Because mechanistic studies of *SETBP1* function have predominantly relied on murine models, it remains unclear whether these epigenetic dependencies are preserved in human HSPCs. Given species-specific differences in hematopoietic regulation and chromatin organization, direct interrogation of *SETBP1*-driven transcriptional control in human progenitors is essential for defining therapeutically relevant vulnerabilities. Here, we investigate the transcriptional and epigenetic mechanisms that sustain *SETBP1*-mutant leukemic programs using primary human HSPC models. By focusing on MENIN-associated KMT2A transcriptional complexes, we test whether *SETBP1*-mutant leukemias exhibit a dependency on MENIN-mediated chromatin regulation and assess the therapeutic relevance of targeting this axis. These studies aim to define actionable epigenetic vulnerabilities in a high-risk molecular subset of myeloid malignancies.

## METHODS

Detailed methods are available in the Supplemental Methods.

### Human CD34^+^ bone marrow stem and progenitor cell culture and lentiviral transduction

Healthy donor bone marrow-derived CD34^+^ HSPCs were obtained from AllCells or StemCell Technologies and cultured in StemSpan SFEM II (StemCell Technologies Cat. #09605) supplemented with StemSpan CD34^+^ Expansion Supplement (StemCell Technologies Cat. #02691) and UM729 (StemCell Technologies Cat. #72332). Codon-optimized human *SETBP1*^*WT*^ and *SETBP1*^*D868N*^ were cloned into the lentiviral pSMAL vector (Addgene #161785). Lentiviral particles were generated in HEK293T-LX cells and concentrated before transduction. CD34^+^ cells were transduced by spininfection, and GFP^+^ cells were purified by flow sorting 48-72 hours after transduction.

### Colony-forming and serial replating assays

GFP^+^ sorted cells were plated into MethoCult H4435 (StemCell Technologies Cat. #4435) semisolid medium in triplicate. Colonies were scored after 14 days using STEMvision^TM^ (StemCell Technologies Cat. #22000). For serial replating assays, cells were harvested, washed, and replated in fresh medium. Colony counts were assessed in ImageJ in a blinded manner.

### CRISPR-Cas9 gene editing

CRISPR-Cas9-mediated knockout of *KMT2A, KMT2B, MEN1*, and *DOT1L* was performed in primary human bone marrow CD34^+^ GFP^+^ sorted Luciferase control or *SETBP1*^*D868N*^ cells using ribonucleoprotein complexes containing SpCas9 protein and chemically synthesized triplet sgRNAs (EditCo, Bio, Inc.). AAVS1 was used as a safe harbor targeting control. Electroporation was performed using a Lonza P3 Primary Cell 4-D Nucleofector X Kit S (Cat. # V4XP-3032) in a Lonza 4D-Nucleofector X Unit (Cat. # AAF-1003X). Knockout efficiency was assessed by Sanger Sequencing and ICE analysis.

### Gene expression studies

RNA was isolated using the Quick-RNA Microprep Kit (Zymo Research, Cat. R1050). cDNA synthesis was performed using a High-Capacity Reverse Transcription kit. Quantitative PCR was performed using TaqMan assays on a QuantStudio 7 instrument. Gene expression was normalized to GUSB and calculated using the 2^-ΔΔCT method. For RNA sequencing, libraries were prepared using the SMART-Seq v4 Plus Kit (Takara, Cat. Nos. R400752& R400753) and sequenced on an Illumina NovaSeq 6000 platform. Reads were aligned to the human genome (GRCh38), and differential gene expression analysis was performed using DESeq2^27^. Gene Set Enrichment Analysis (GSEA)^28,29^ was performed using curated (C2; CGP and Hallmark) gene sets from the Molecular Signatures Database (MSigDB)^30^.

### CUT&Tag

CUT&Tag was performed with antibodies against SETBP1, MENIN, KMT2A, H3K4me3, and H3K27ac, with IgG serving as a control. Libraries were sequenced on an Illumina NextSeq 2000 platform. Reads were aligned to GRCh38, peaks were identified using GoPeaks, and downstream analysis was performed using BedTools and deepTools. Integration of chromatin occupancy and transcriptional profiling was performed using a Python 3 implementation of BETA^31^.

### Flow Cytometry

Cells were incubated with fluorochrome-conjugated antibodies (CD117, CD11b, CD14, CD15, CD16, CD66b, HLA-DR) and assessed on LSR Fortessa (BD Biosciences). Further analysis was performed with FlowJo software (Tree Star). Population gating was performed according to an unstained sample.

### Menin inhibitor studies

Ziftomenib, Revumenib and Bleximenib were evaluated in *SETBP1-*mutant cell lines, engineered CD34^+^ HSPCs and primary *SETBP1*-mutant patient samples. Effects on proliferation, colony formation, differentiation, gene expression and chromatin occupancy were assessed as described above.

### Data presentation

All graphs or figures were made using GraphPad Prism or BioRender. Figures were assembled in Affinity Designer. Statistical significance was evaluated using GraphPad Prism. Data distribution was assumed to be normal. Data are presented as mean (+/-) standard error of the mean. Statistical tests are indicated in the figure legends. P-values < 0.05 were considered statistically significant.

## RESULTS

### Mutant *SETBP1* promotes self-renewal and myeloid progenitor expansion through activation of a *HOXA/MEIS1*-driven leukemogenic transcriptional program

Previously, the functional consequences of *SETBP1* mutations have been largely studied in murine models, limiting clinical relevance. To understand the role of mutant *SETBP1* in human hematopoiesis, we transduced primary human bone marrow CD34^+^ HSPCs with a control lentiviral vector (*psmal-Luc*), *psmal-SETBP1*^*WT*^, or *psmal-SETBP1*^*D868N*^ (**Figure 1A**). Expression of *SETBP1*^*D868N*^ in CD34^+^ cells significantly enhanced cell proliferation compared with both control and *SETBP1*^*WT*^ cells (**Figure 1B**). To determine whether mutant *SETBP1* enhances progenitor self-renewal, we performed colony-forming unit (CFU) serial replating assays. *SETBP1*^*D868N*^ markedly increased colony numbers in the second plating, indicating enhanced progenitor self-renewal potential (**Figure 1C-E**). In addition, CFU analysis revealed that *SETBP1*-mutant cells had increased granulomonocytic lineage output upon replating (**Figure S1A, B**).

**Figure 1.**
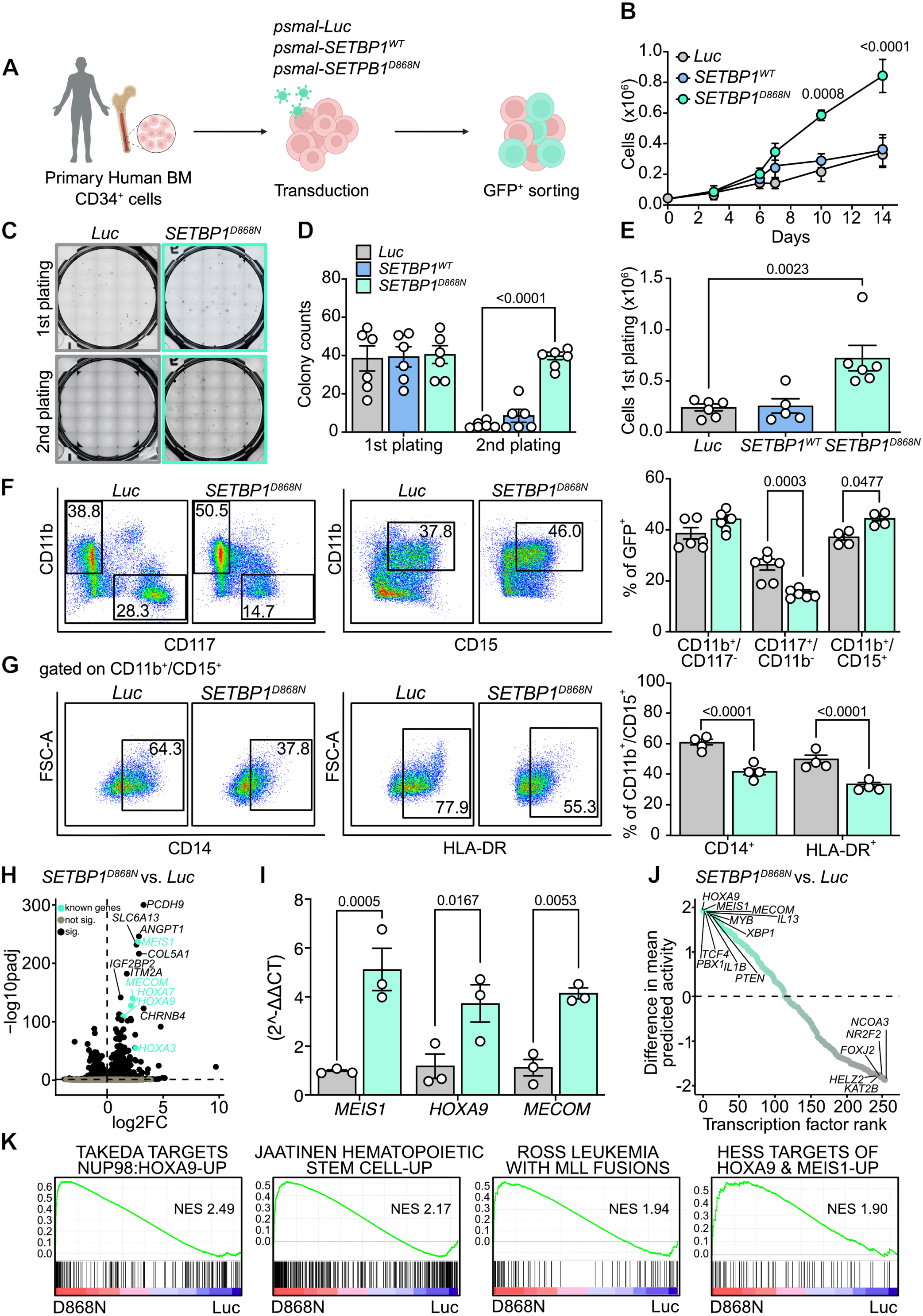
*SETBP1*-mutant human CD34^+^ cells activate a HOXA/MEIS1 leukemogenic program A. Transduction of primary human bone marrow (BM) CD34^+^ cells with wildtype (*psmal-SETBP1*^*WT*^), mutant (*psmal-SETBP1*^*D868N*^) or control (*psmal-Luc*) constructs. GFP-positive cells were sorted for functional downstream analyses. **B**. Growth curve of cells after sorting over 14 days in liquid culture (n=3 for *Luc*, n=4 for *SETBP1*^*D868N*^, *and* n=3 for *psmal-SETBP1*^*WT*^) **C**. Transduced cells from (A) were plated in methylcellulose-based colony-forming medium (2500 cells/well). Representative CFU assay images of *Luc* and *SETBP1*^*D868N*^-expressing cells are shown. **D**. Quantification of CFU assays. The number of colonies from (B) was counted in biological triplicate wells after 14 days. **E**. Quantification of isolated cells from CFU assays after 1^st^ plating. **F**. Representative flow cytometry images and quantification of *Luc* and *SETBP1*^*D868N*^-expressing cells cultured for 13 days after sorting are shown. CD11b^+^/CD117^-^, CD117^+^/CD11b^-^, and CD11b^+^/CD15^+^ cells were gated on GFP-positive cells. **G**. Representative flow cytometry images and quantification of *Luc* and *SETBP1*^*D868N*^-expressing cells cultured for 13 days after sorting are shown. CD14^+^ and HLA-DR^+^ cells were gated on CD11b^+^/CD15^+^ cells. **H**. RNA-sequencing analysis of *SETBP1*^*D868N*^-transduced versus control-transduced primary human CD34^+^ cells (n=5 per group) seven days after sorting. Volcano plot of differentially expressed genes in each condition. Significantly upregulated or downregulated genes (log fold change > 0 or <0, black). Among the significantly upregulated genes (padj < 0.05) those highlighted in aquamarine correspond to known SETBP1-target genes; p-values were calculated using a Wald test with Benjamini-Hochberg false discovery rate (FDR) correction for multiple comparisons, as implemented in DESeq2. Due to numeric underflow in statistical calculations, some genes were assigned adjusted p-values of 0, resulting in infinite -log10(padj) values. For visualization, these values were capped at 300. **I**. qRT-PCR analysis of *HOXA9, MEIS1*, and *MECOM* mRNA transcripts from liquid culture harvested at 14 days after sorting. Expression levels were normalized to the *GUSB* housekeeping gene. **J**. Transcription factors ranked by the difference in their mean predicted activity in *SETBP1*^*D868N*^ versus control cells. Predicted activity scores were determined using Priori, and points are colored by the magnitude and direction of the difference. Top-ranked TFs with higher mean predicted activity in the *SETBP1*^*D868N*^ condition are in turquoise, and TFs with higher mean predicted activity in the control are in gray. **K**. Gene Set Enrichment Analysis (GSEA) of *SETBP1*^*D868N*^ and control conditions using C2: CGP and Hallmark gene sets (q-value < 0.05). Positive normalized enrichment scores (NES) for the top gene sets enriched in *SETBP1*^*D868N*^*-*expressing cells. If not otherwise described, statistical analyses were performed with 2-way ANOVA with Sidak’s multiple comparison test. Error bars represent the standard error of the mean (SEM) for B, D, E, F, G and I. A paired t-test was used for E.

To further determine how *SETBP1* mutations affect myeloid progenitor expansion and maturation, we performed flow cytometric immunophenotyping of *SETBP1*-mutant cells. Compared with controls, *SETBP1*^*D868N*^ HSPCs displayed a less stem-like, more mature myeloid progenitor phenotype, characterized by decreased CD117 and increased CD11b surface marker expression (**Figure 1F**). However, these cells remained largely within the progenitor compartment and did not undergo terminal differentiation. Consistent with impaired myeloid maturation, *SETBP1*^*D868N*^ expression reduced the proportion of cells expressing markers of differentiated monocytic and granulocytic cells. Specifically, *SETBP1*-mutant cells exhibited fewer CD14^+^/HLA-DR^+^ monocytic cells and reduced expression of the granulocytic marker CD66b (**Figure 1G, Figure S1C**). Furthermore, expression of the terminal neutrophil marker CD16 remained low overall but was further reduced in *SETBP1*-mutant cells compared to controls (**Figure S1D**).

To define transcriptional programs associated with mutant *SETBP1*, we performed RNA-sequencing of control and *SETBP1*^*D868N*^-expressing human CD34^+^ HSPCs. Differential gene expression analysis revealed marked upregulation of canonical stemness- and leukemia-associated genes, including *MEIS1, HOXA* cluster genes, and *MECOM* (**Figure 1H, Table S1**). We validated increased expression of these key regulators by qPCR, confirming the RNA-seq results and showing that *SETBP1*^*D868N*^ promotes acquisition of a stem cell-associated transcriptional program (**Figure 1I**). To identify regulatory drivers of this expression profile, we applied Priori-based transcription factor activity analysis^32^. Among the highest-ranked predicted regulators were HOXA9, MEIS1, MECOM and MYB, all of which have established roles in hematopoiesis, stem cell maintenance and leukemogenesis **(Figure 1J, Table S2)**. Consistent with these predictions, the transcriptional landscape of *SETBP1*^*D868N*^ cells was characterized by activation of HOX- and MEIS-associated regulatory programs. Gene set enrichment analysis (GSEA) revealed enrichment of NUP98-HOXA9 fusion targets and MLL fusion leukemia signatures in *SETBP1*^*D868N*^-expressing HSPCs, while these cells also show upregulation of HOXA and MEIS1 targets and hematopoietic stem cell genes (**Figure 1K)**.

Collectively, these findings demonstrate that *SETBP1*^*D868N*^ reprograms human HSPCs toward a stem-like leukemic state characterized by enhanced proliferation and self-renewal, activation of *HOXA/MEIS1*-associated transcriptional programs, and accumulation of immature myeloid progenitor-like cells.

### Mutant *SETBP1* establishes a KMT2A-MENIN-associated chromatin program at leukemogenic regulatory elements

Having identified that mutant *SETBP1* alters transcriptional programs toward a *HOXA/MEIS1*-driven leukemogenic state, similar to KMT2A-rearranged leukemias, we next set out to investigate the chromatin mechanisms underlying enhanced self-renewal and activation of stem cell-associated genes. To this end, we performed CUT&Tag profiling in control and *SETBP1*^*D868N*^ human HSPCs. Expression of *SETBP1*^*D868N*^ resulted in increased chromatin occupancy of SETBP1, relative to control cells (**Figure 2A**). Notably, in *SETBP1*-mutant HSPCs, we observed a global increase in chromatin-bound KMT2A, while MENIN occupancy remained largely unchanged (**Figure 2B, C**). To further define the genomic context of SETBP1-associated chromatin regulation, we examined the distribution of SETBP1, KMT2A, and MENIN occupancy across genomic features. SETBP1^D868N^ binding was enriched at promoter regions, whereas KMT2A and MENIN individually displayed a broader genomic distribution. Sites co-occupied by KMT2A and MENIN alone were predominantly localized to distal intergenic regions. In contrast, loci jointly occupied by SETBP1^D868N^, KMT2A, and MENIN showed a marked shift toward promoters (55%) and a smaller fraction occurring at intergenic regions (**Figure S2A**). These findings suggested that mutant SETBP1 promotes establishment of a promoter-focused KMT2A-MENIN chromatin program at shared target loci.

**Figure 2.**
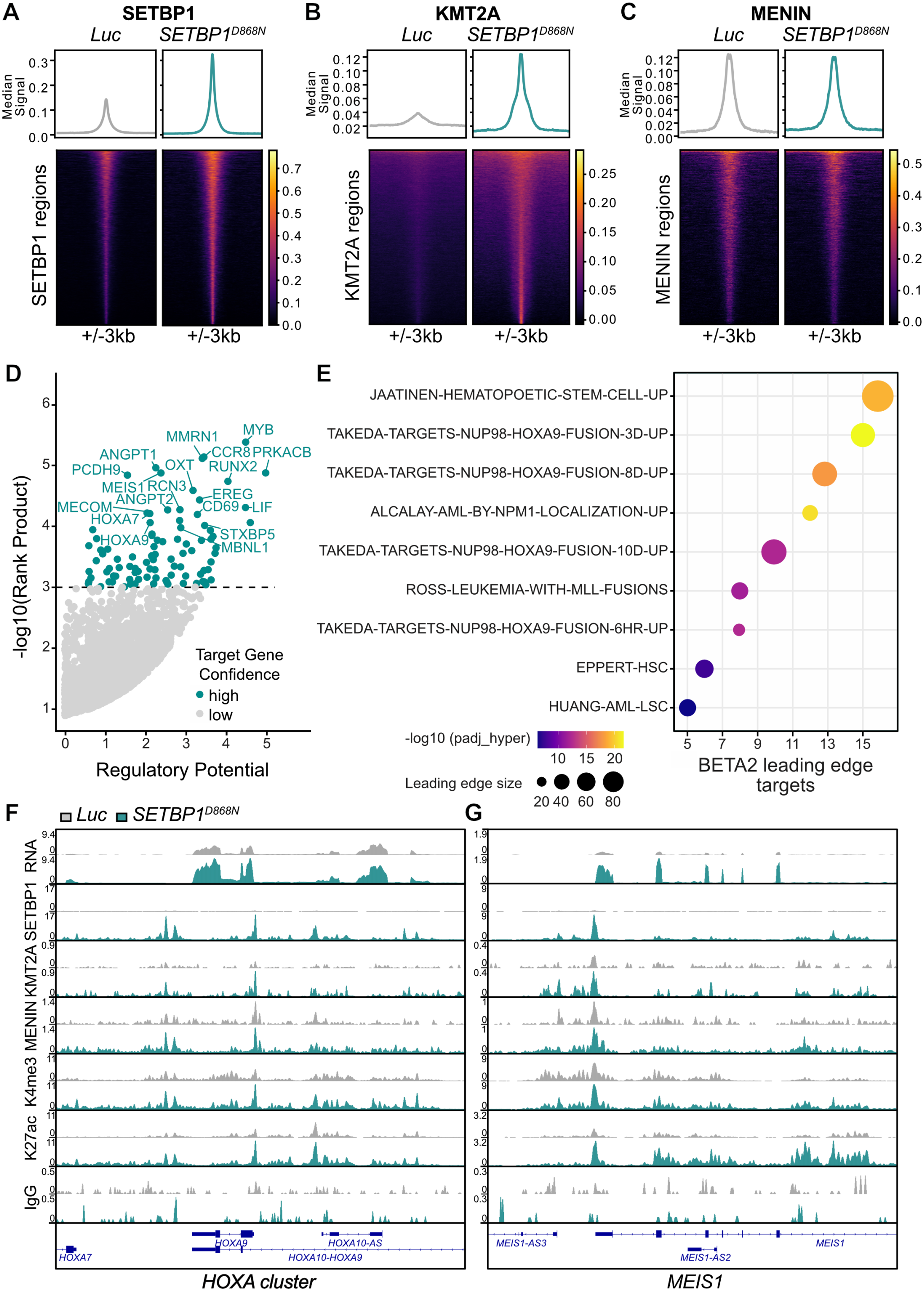
High correlation between SETBP1-bound regions with KMT2A binding and its cofactor MENIN. A-C. CUT&Tag for (A) SETBP1, (B) KMT2A, and (C) MENIN in primary human bone marrow CD34^+^ cells with mutant (*psmal-SETBP1*^*D868N*^) or control (*psmal-Luc*) constructs in biological triplicate. Heatmaps and median signal profiles of normalized SETBP1, KMT2A, or MENIN CUT&Tag signal in control (*Luc*) and *SETBP1*^*D868N*^ conditions (n=3 per group and condition, except Luc-MENIN is n=2). The signal is shown at peak centers +/-3kb. Each row in the heatmap represents one binding region. **D**. Predicted SETBP1 regulome identified by integrating CUT&Tag occupancy with differential gene expression using the BETA algorithm. Genes nominated by BETA as SETBP1 targets and transcriptionally upregulated in *SETBP1*^*D868N*^ HSPCs are highlighted in teal. **E**. GSEA leading-edge analysis of BETA-nominated SETBP1 target genes. Overlap between BETA-nominated SETBP1 targets and leading-edge genes of each gene set was assessed using a one-sided hypergeometric test with Benjamini-Hochberg (BH) multiple-testing correction. Dot color represents -log10 (BH-adjusted p-value), and dot size represents the number of genes in each pathway’s leading edge. Only GSEA pathways with positive NES (more enriched in the D868N condition) were included in the analysis. **F-G**. Merged IGV tracks for RNA-seq expression data (n=5 per group), SETBP1 (n=3 per group), KMT2A (n=3 per group), MENIN (n=3 for *SETBP1*^*D868N*^, n=2 for *Luc*), H3K4me3 (n=3 per group), and H3K27ac (n=3 per group) are shown at the (F) *HOXA cluster* or (G) *MEIS1* locus. IgG serves as a control.

To identify genes directly regulated by this chromatin program, we integrated differential gene expression and SETBP1 chromatin occupancy using Binding and Expression Target Analysis (BETA^31^). BETA2^33^ is a Python 3 implementation that integrates CUT&Tag data with differential gene expression to infer direct target genes. This analysis identified a high-confidence predicted SETBP1 regulome comprising genes that were both transcriptionally upregulated and directly bound by *SETBP1*^*D868N*^. In total, 94 genes met these criteria, including established key leukemogenic regulators such as HOXA9, MEIS1, MYB, and RUNX2, supporting the robustness of this approach (**Figure 2D, Table S3**). To define the biological programs represented by the SETBP1 regulome, we examined its overlap with leading-edge genes from enriched GSEA gene sets. This revealed that the SETBP1-regulated genes significantly overlap with genes driving transcriptional programs associated with *KMT2A*-rearranged leukemia, *NPM1*-mutant AML, *NUP98-HOXA9*-driven leukemia, and hematopoietic and leukemia stem cell self-renewal signatures (**Figure 2E, Table S4**).

Given the prominent enrichment of KMT2A-associated leukemogenic programs within the SETBP1 regulome, we next asked whether mutant *SETBP1* was associated with an altered chromatin state at these target loci. CUT&Tag profiling of active chromatin marks H3K4me3 and H3K27ac revealed that SETBP1-regulated loci, such as the *HOXA* cluster, *MEIS1* and *MYB*, showed increased H3K4me3 and H3K27ac in *SETBP1*^*D868N*^ HSPCs relative to controls **(Figure 2F, G, Figure S2B**). These loci also displayed increased SETBP1, KMT2A and MENIN occupancy, linking promoter-focused co-occupancy with establishment of an active chromatin state.

Together, these findings support a model in which mutant SETBP1 redistributes KMT2A-MENIN occupancy toward promoter-associated leukemogenic targets, where it facilitates acquisition of a transcriptionally permissive chromatin landscape and activates oncogenic gene expression.

### KMT2A and MENIN are selective molecular dependencies in *SETBP1-*mutant HSPCs

Having identified increased KMT2A occupancy at SETBP1 target loci, we next sought to determine whether components of the KMT2A-MENIN complex are functionally required for SETBP1-driven leukemogenesis. In addition to occupying KMT2A-associated target genes, SETBP1 has been reported to interact with multiple members of the KMT2 methyltransferase complex, including KMT2A, KMT2B and DOT1L^18^ (**Figure 3A**). Consistent with a functional relationship between SETBP1 and the KMT2A-MENIN complex, SETBP1 and MENIN were found to co-immunoprecipitate in *SETBP1*^*D868N*^ cells (**Figure S3A)**.

**Figure 3.**
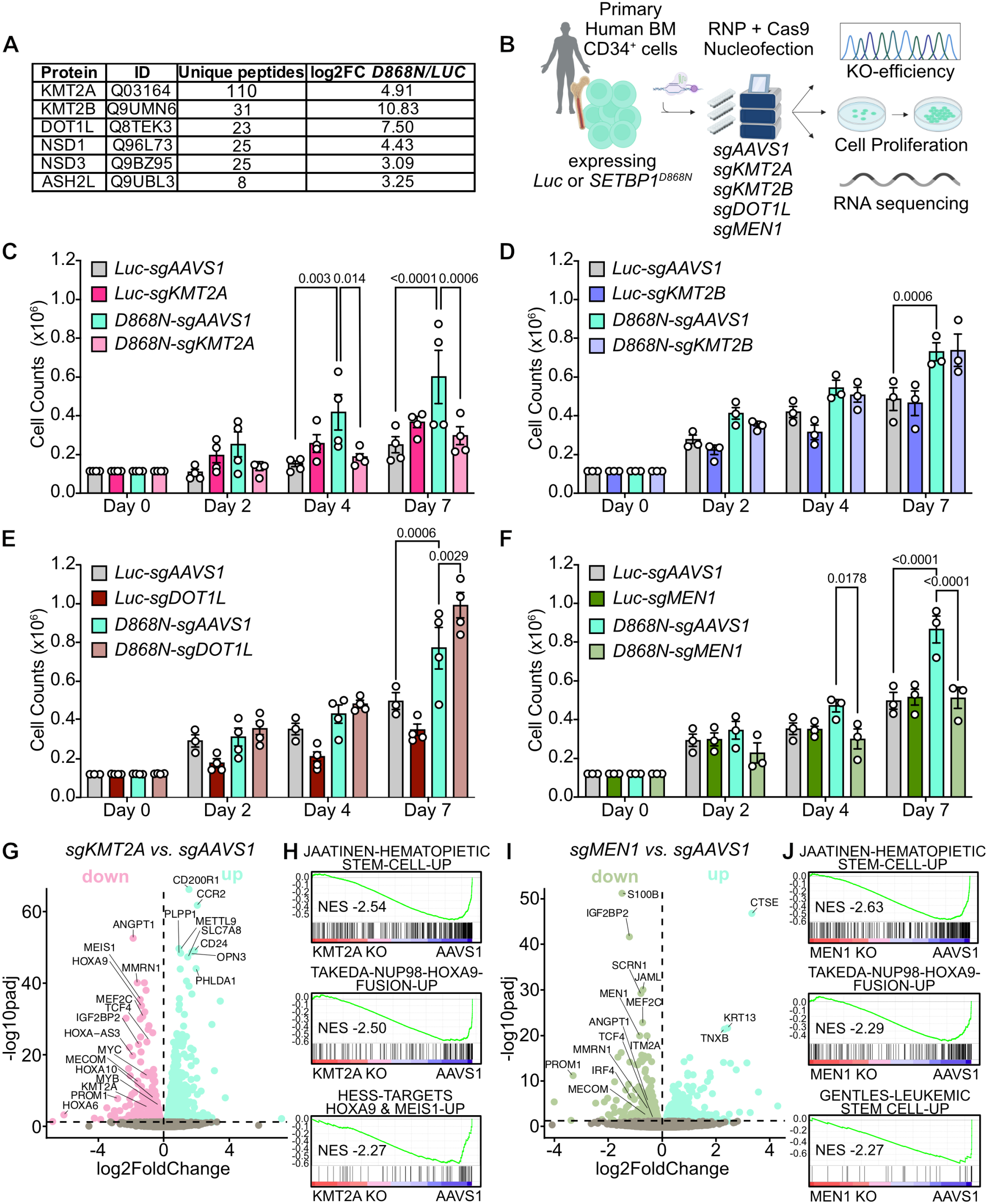
KMT2A and MENIN are dependencies in *SETBP1*^*D868N*^ hematopoietic progenitors. **A**. Unique peptides identified in BioID in BirA control or BirA-SETBP1^D868N^ transduced K562 cells. Relevant HMTs identified in the screen are shown. **B**. Schematic for CRISPR/Cas9 modification of primary human CD34^+^ HSPCs. **C-F**. Growth of control or *SETBP1*^*D868N*^ HSPCs modified with a non-targeting control triplet guide (*sgAAVS1*) or guides targeting **(A)** *KMT2A*, **(B)** *KMT2B*, **(C)** *DOT1L*, or **(D)** *MEN1* over 7 days after nucleofection **E-F**. RNA-seq analysis of *SETBP1*^*D868N*^-transduced primary human CD34^+^ cells following **(E)** KMT2A (sgKMT2A) or **(F)** MEN1 (sgMEN1) knockout after 7 days. AAVS1 (sgAAVS1) served as a control. **G-J**. Volcano plot of differentially expressed genes in each condition. Significantly upregulated genes are shown in turquoise (log fold change > 0, padj <0.05); significantly downregulated genes are shown in pink for **(G)** sgKMT2A (n=3) or green for **(I)** sgMEN1 (n=4) (log fold change <0, padj < 0.05). **(H, J)** Gene set enrichment analysis (GSEA) was performed using C2: CGP and Hallmark gene sets. Negative normalized enrichment scores (NES) indicate reduced enrichment of the indicated gene set following knockout. If not otherwise described, statistical analyses were performed with 2-way ANOVA with Sidak’s multiple comparison test. Error bars represent standard error of the mean (SEM) for C, D, E, and F.

To evaluate their functional requirement, we performed high-efficiency CRISPR-Cas9-mediated knockout studies using a triplet-guide strategy, achieving 60-100% knockout efficiency (**Figure 3B, Figure S3B-F**). Loss of KMT2A markedly impaired cell growth in *SETBP1*-mutant HSPCs (**Figure 3C**). In contrast, disruption of KMT2B did not affect cellular proliferation, while loss of DOT1L unexpectedly led to modestly increased cell growth in *SETBP1*-mutant HSPCs (**Figure 3D, E**). Importantly, deletion of MEN1 phenocopied the effects of KMT2A loss, resulting in a profound reduction in growth of *SETBP1*-mutant cells (**Figure 3F**). Notably, knockout of KMT2A and MEN1 had little effect on control HSPCs, suggesting the potential for a therapeutic window. Together, these data identify the KMT2A-MENIN complex as a specific molecular dependency in *SETBP1*-mutant cells.

To determine how KMT2A or MENIN support *SETBP1*-driven leukemogenic programs, we performed RNA-sequencing seven days following CRISPR-mediated gene disruption of KMT2A or MEN1. Loss of KMT2A significantly reduced the expression of canonical *SETBP1*^*D868N*^ target genes, including the *HOXA* cluster, *MEIS1, MECOM, MYB*, and *MYC*, as well as genes associated with leukemia stem cell maintenance and hematopoietic progenitor identity, including *MMRN1, PCHD9, ANGP1, MEF2C* and *PROM1* (**Figure 3G, Table S5**).

Similarly, disruption of MEN1 suppressed expression of key leukemogenic regulators, including *MEF2C, ANGPT1, MMRN1, MECOM, PROM1, TCF4*, and I*RF4* (**Figure 3I, Table S6**). GSEA revealed broad suppression of stemness-associated and leukemogenic transcriptional programs following loss of either KMT2A or MENIN, including hematopoietic stem cell signatures, *HOXA9* and *MEIS1* target genes, and *NUP98:HOXA9*-fusion signatures (**Figure 3H, J**). Consistent with these findings, EnrichR analysis using the KEGG 2026 database and cell-type specific enrichment using CellMarker 2024 database demonstrated significant loss of hematopoietic stem and progenitor cell-associated programs in both perturbation settings (**Figure S4A-D**).

Collectively, these findings identify KMT2A and MENIN as critical dependencies in *SETBP1*-mutant cells. Genetic disruption of either KMT2A or MEN1 impaired proliferation and suppressed expression of stemness-associated and leukemogenic transcriptional programs driven by *SETBP1*^*D868N*^, identifying the KMT2A-MENIN complex as a key therapeutic vulnerability in *SETBP1*-mutant leukemia.

### Ziftomenib disrupts *SETBP1*-driven self-renewal and promotes hematopoietic differentiation

Having identified the KMT2A-MENIN complex as a critical dependency in *SETBP1*-mutant cells, we next investigated whether pharmacologic inhibition of the KMT2A-MENIN interaction could disrupt mutant *SETBP1*-driven leukemogenic programs. Treatment of the SETBP*1*-mutant MOLM-20 cell line with the MENIN inhibitor ziftomenib only modestly reduced cell growth but significantly suppressed the expression of canonical SETBP1 target genes, including *HOXA9, MEIS1*, and *MECOM* (**Figure 4A, B**). Similarly, treatment of primary *SETBP1*^*D868N*^ HSPCs for seven days had minimal effects on short-term proliferation (**Figure 4C**).

**Figure 4.**
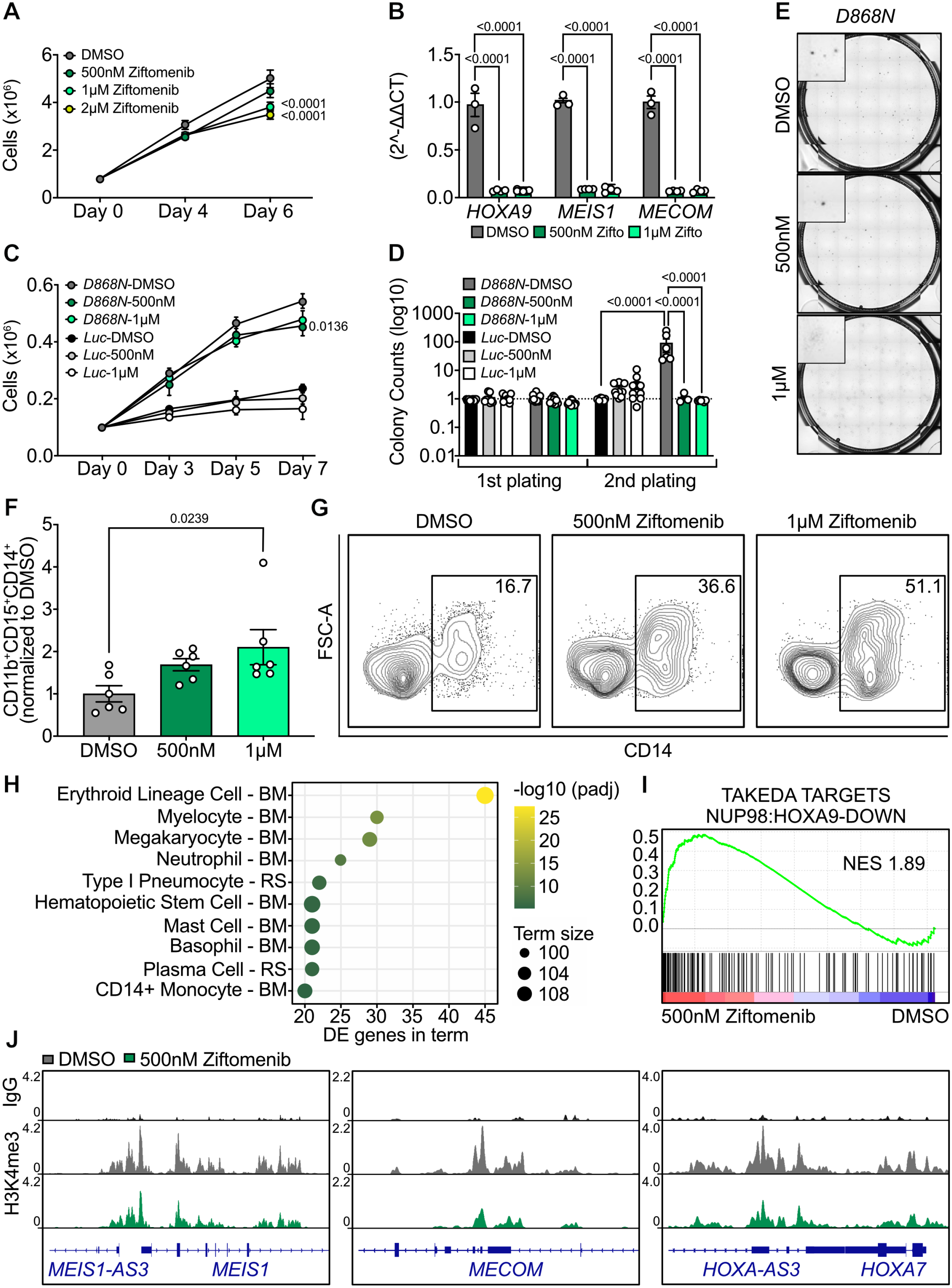
Inhibition with ziftomenib selectively impairs clonogenic growth, suppresses oncogenic gene expression and disrupts KMT2A-associated chromatin occupancy. **A**. Growth curve of SETBP1-mutant MOLM-20 cells treated with DMSO, 500nM, 1μM or 2μM of ziftomenib in liquid culture for 6 days. **B**. qRT-PCR analysis of *HOXA9, MEIS1*, and *MECOM* mRNA transcripts after six days of treatment of MOLM-20 cells with 500nM or 1μM of ziftomenib or DMSO. Expression levels were normalized to the *GUSB* housekeeping gene and to DMSO. **C**. Growth curve of control or *SETBP1*^*D868N*^ HSPCs treated with DMSO, 500nM or 1μM of ziftomenib in liquid culture for 7 days. **D**. Control or *SETBP1*^*D868N*^ HSPCs plated in methylcellulose-based colony-forming medium (2500 cells/well) and treated with DMSO, 500nM or 1μM of ziftomenib. After 14 days, cells were replated at a concentration of 25,000 cells/well. **E**. Representative CFU assay images of *SETBP1*^*D868N*^-expressing cells after 14 days of ziftomenib treatment are shown. **F**. Quantification of flow cytometric analysis from isolated *SETBP1*^*D868N*^ HSPCs from CFU assays after 1^st^ plating. Ordinary 1-way ANOVA with Dunnett’s multiple comparisons test. Percentages were normalized to DMSO. CD14^+^ cells were gated on GFP^+/^CD11b^+^/CD15^+^ cells. **G**. Representative flow cytometry images of *SETBP1*^*D868N*^ HSPCs treated with DMSO, 500 nM, or 1μM ziftomenib are shown. **H**. EnrichR analysis of differentially expressed genes in *SETBP1*^*D868N*^ HSPCs (500nM ziftomenib, n=3; DMSO, n=4) following six days of ziftomenib treatment using the HuBMAP ASCTplusB augmented 2022 library. The top 10 enriched cell-type signatures are shown. The x-axis indicates the number of differentially expressed (DE) genes overlapping each signature; dot color indicates the adjusted p-value, and dot size indicates number of genes in each signature. BM, bone marrow; RS, respiratory system. **I**. Gene set enrichment analysis (GSEA) was performed using C2: CGP and Hallmark gene sets. Positive normalized enrichment scores (NES) indicate increased enrichment of the indicated gene set following ziftomenib treatment. **J**. Merged tracks for H3K4me3 (n=3 per group) following three days of treatment of *SETBP1*^*D868N*^ HSPCs with ziftomenib are shown at the *HOXA cluster, MEIS1 and MECOM* loci. IgG serves as a control. If not otherwise described, statistical analyses were performed with 2-way ANOVA with Sidak’s multiple comparison test. Error bars represent standard error of the mean (SEM) for A, B, C, D and F.

Given the effects on stemness-associated gene expression, we next examined whether MENIN inhibition affected the enhanced self-renewal capacity conferred by mutant *SETBP1*. Serial CFU assays revealed that ziftomenib abolished the enhanced secondary replating capacity of *SETBP1*^*D868N*^-expressing cells, whereas colony formation by control cells was largely unaffected (**Figure 4D**). Remaining colonies derived from *SETBP1*^*D868N*^ cells displayed a more differentiated morphology, consistent with previous reports demonstrating that MENIN inhibition promotes myeloid differentiation (**Figure 4E**). Accordingly, ziftomenib treatment increased the proportion of CD11b^+^/CD15^+^/CD14^+^ expressing cells (**Figure 4F, G**).

To further define the differentiation response, we performed RNA-sequencing of *SETBP1*^*D868N*^ HSPCs following six days of ziftomenib treatment in liquid culture. Differential gene expression analysis followed by EnrichR revealed enrichment of hematopoietic differentiation signatures including erythroid cells, myelocytes, megakaryocytes, basophils, neutrophils and CD14^+^ monocytes-associated signatures (**Figure 4H, Table S7**). Among the most highly upregulated differentiation-associated genes were *TRIM29, EPHB4, RHCE, TNFRSF19, HBA2, PKLR, HBA1, MYH10, ITGB4, ESPN, KLF1, FAM178B, ALAS2, SLC6A19, CA2, TRIB2*, and *MYO10* (**Figure S5A**). In contrast, DMSO-treated *SETBP1*^*D868N*^ HSPCs maintained higher expression of progenitor- and leukemia-associated genes, including *CD34, MEF2C*, and *IGF2BP2* (**Figure S5A**). GSEA further demonstrated depletion of the NUP98:HOXA9 fusion signature and enrichment of heme metabolism and mature hematopoietic cell programs following ziftomenib treatment of *SETBP1*^*D868N*^ HSPCs, while pathways associated with undifferentiated cancers were enriched in the DMSO control condition (**Figure 4I, Figure S5B-D**).

To investigate the chromatin consequences of MENIN inhibition, we performed CUT&Tag profiling following three days of ziftomenib treatment. Although technical limitations precluded direct profiling of SETBP1, MENIN or KMT2A, ziftomenib reduced H3K4me3 occupancy at key leukemogenic loci including *MEIS1, MECOM* and *HOXA* cluster, indicating loss of transcriptionally permissive chromatin at critical SETBP1 target genes (**Figure 4J**).

Collectively, these findings demonstrate that pharmacological inhibition of the KMT2A-MENIN interaction disrupts the chromatin and transcriptional programs that sustain mutant *SETBP1*-driven leukemogenesis. Despite only modest effects on short-term proliferation, ziftomenib reduced expression of key leukemogenic regulators, reduced H3K4me3 at SETBP1 target loci, eliminated the enhanced self-renewal and promoted myeloid differentiation of *SETBP1*-mutant cells. These results support a model in which MENIN inhibition acts primarily as a differentiation-directed therapeutic strategy that dismantles leukemic stem cell programs rather than inducing direct cytotoxicity, highlighting the KMT2A-MENIN complex as a therapeutically actionable dependency in *SETBP1*-mutant leukemia.

### *SETBP1*-mutant patient leukemias are sensitive to pharmacological KMT2A-MENIN inhibition

To translate our findings to clinically relevant models, we evaluated the effects of MENIN inhibition in primary patient samples harboring recurrent *SETBP1* mutations. Six independent *SETBP1*-mutant patient samples were treated with three clinically relevant MENIN inhibitors: ziftomenib, revumenib, and bleximenib. Treatment with all three MENIN inhibitors significantly reduced clonogenic output across *SETBP1*-mutant samples (**Figure 5A-B**). Notably, the reduction in colony formation was primarily driven by impaired proliferative expansion within colonies rather than a substantial decrease in the total number of colonies formed (**Figure 5C**). In contrast, treatment of normal CD34^+^ HSPCs resulted in only modest changes in colony formation, suggesting selectivity for *SETBP1*-mutant cells (**Figure S5E**).

**Figure 5.**
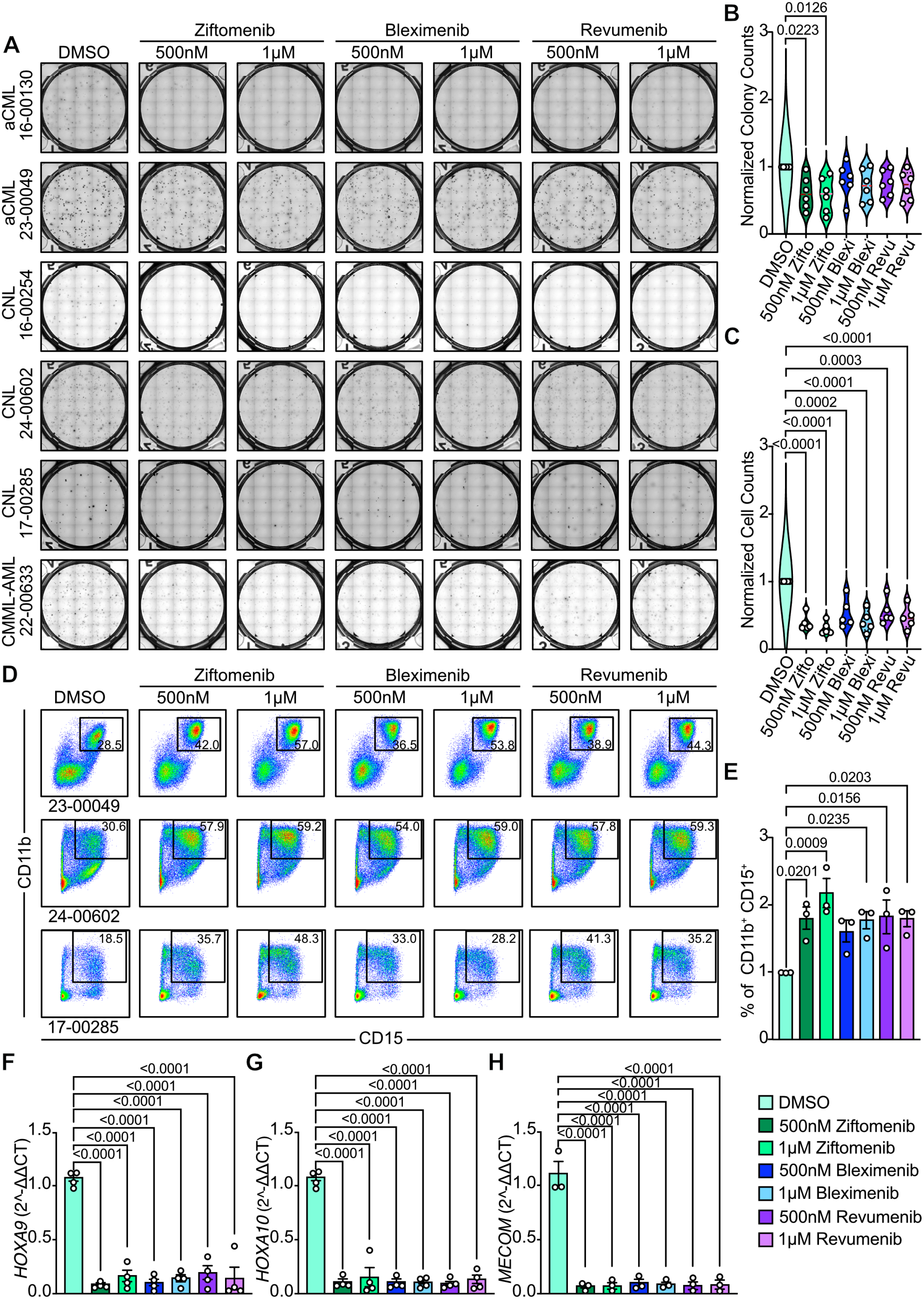
Menin inhibition is effective in samples from patients with *SETBP1*-mutant leukemia. **A**. Representative images from CFU assays for six different patient samples that were treated with DMSO, 500 nM, or 1µM with three different menin inhibitors: ziftomenib, bleximenib, and revumenib for 14 days. **B**. Normalized colony counts after 14 days of treatment with ziftomenib, bleximenib or revumenib **C**. Normalized cell counts from colony-forming unit assays after 14 days of treatment with ziftomenib, bleximenib or revumenib. **D**. Representative flow cytometry analysis of CD11b and CD15 surface expression harvested from colonies at 14 days. **E**. Quantification of flow cytometry surface markers CD11b^+^/CD15^+^ cells normalized to DMSO. Each replicate represents a biologically independent sample from a distinct patient. **F**. qRT-PCR analysis of *HOXA9, HOX10*, and *MECOM* mRNA transcripts from colonies harvested at 14 days. Each replicate represents a biologically independent sample from a distinct patient. Expression levels were normalized to the *GUSB* housekeeping gene and DMSO. Statistical analyses were performed with ordinary 1-way ANOVA with Dunnett’s multiple comparisons test. Error bars represent standard error of the mean (SEM) for B, C, E, F, G, and H.

Given the effects of MENIN inhibition on clonogenic growth, we next investigated whether these changes were accompanied by altered differentiation. Flow cytometric analysis of colonies derived from *SETBP1*-mutant patient samples treated with MENIN inhibitors showed increased expression of myeloid differentiation markers CD11b and CD15, consistent with enhanced myeloid maturation (**Figure 5D, E**). These phenotypic changes were accompanied by suppression of key leukemogenic programs. Expression of canonical *SETBP1* target genes, including *HOXA9, HOXA10* and *MECOM*, was significantly reduced following MENIN inhibition (**Figure 5F-H)**. While *MEIS1* expression was decreased in one patient sample, this effect was not consistently observed across all samples, potentially due to the low baseline expression in these specimens. (**Figure S5F**). Similarly, *MYC* expression showed substantial inter-patient variability and was not significantly altered overall (**Figure S5G**).

Collectively, these findings demonstrate that *SETBP1*-mutant patient leukemias remain dependent on KMT2A-MENIN interaction to maintain clonogenic growth and leukemogenic transcriptional programs. Pharmacological MENIN inhibition suppresses expression of key *SETBP1* target genes while promoting myeloid differentiation and limiting progenitor expansion. The consistent activity of multiple MENIN inhibitors across genetically distinct *SETBP1*-mutant patient samples validate the KMT2A-MENIN interaction as a therapeutically actionable vulnerability and provides a strong rationale for the clinical evaluation of MENIN-targeted therapies in *SETBP1*-mutant myeloid malignancies.

## DISCUSSION

*SETBP1* mutations are recurrent in myeloid malignancies and are consistently associated with aggressive disease biology and poor clinical outcomes. Although previous studies established a role for mutant *SETBP1* in leukemogenesis, much of our mechanistic understanding has been derived from murine models^17,18,21^. Here we demonstrate that mutant *SETBP1* reprograms human HSPCs toward a self-renewing leukemogenic state characterized by activation of HOXA/MEIS1 transcriptional programs and enhanced clonogenic potential. Importantly, we identify the KMT2A-MENIN complex as a critical mediator of these effects and uncover a therapeutically actionable dependency in *SETBP1*-mutant leukemia.

Consistent with prior murine studies, *SETBP1*^*D868N*^ similarly enhanced proliferation and self-renewal of primary human CD34^+^ HSPCs (**Figure 1**)^1,12,18^. Moreover, mutant *SETBP1* promoted expansion of immature myeloid progenitors accompanied by activation of canonical leukemogenic regulators, including *HOXA9, MEIS1*, and *MECOM*^19,34–36^. These transcriptional changes closely resemble those observed in *KMT2A*-rearranged, *NPM1*-mutant, and *NUP98-HOXA9*-driven leukemias, suggesting that distinct genetic lesions may converge on common stemness-associated transcriptional networks^37–41^.

A central finding of our studies is that mutant *SETBP1* remodels chromatin at key leukemogenic loci through engagement of the KMT2A-MENIN complex (**Figure 2**). Integration of chromatin occupancy and transcriptional profiling predicted a high-confidence SETBP1 regulome enriched for leukemia stem cell and KMT2A-associated gene signatures. *SETBP1*-mutant cells displayed increased KMT2A occupancy together with enrichment of active chromatin marks, including H3K4me3 and H4K27ac at *HOXA* cluster genes, *MEIS1*, and other direct SETBP1 targets. Although global MENIN occupancy was not increased, MENIN remained localized at these loci, suggesting that mutant *SETBP1* may promote the stabilization or activation of KMT2A-MENIN complexes rather than recruiting MENIN *de novo*. Together, these findings suggest that mutant *SETBP1* establishes a KMT2A-MENIN chromatin program that sustains leukemogenic transcriptional programs.

Our functional studies further demonstrate that the *SETBP1*-mutant transcriptional program is critically dependent on KMT2A and MENIN (**Figure 3**). Genetic disruption of either KMT2A or MEN1 impaired proliferation of *SETBP1*-mutant cells and suppressed expression of key leukemogenic regulators, extending previous studies in murine systems implicating KMT2A as a critical mediator of *SETBP1-* driven transformation in primary human HPSCs^17,21^. These findings place *SETBP1*-mutant cells within a broader group of leukemias that depend on MENIN-KMT2A-mediated maintenance of *HOXA/MEIS1* transcriptional programs^24,37^. Notably, loss of KMT2A or MEN1 had minimal effects on control HSPCs, supporting a therapeutic window.

The clinical relevance of these findings is underscored by our studies using pharmacological inhibitors targeting the KMT2A-MENIN interaction^26,42,43^. In *SETBP1*-mutant HSPCs, ziftomenib had modest effects on short-term proliferation but abolished the enhanced serial replating and promoted myeloid differentiation (**Figure 4**). Transcriptomic profiling further supported a broader differentiation response, suggesting that MENIN inhibition disrupts the progenitor-like state maintained by mutant SETBP1 and permits engagement of multiple hematopoietic differentiation programs, rather than directing differentiation toward a single lineage. These phenotypic changes were accompanied by reduced H3K4me3 at *MEIS1, MECOM* and *HOXA*-cluster loci, supporting a model in which disruption of permissive chromatin at key SETBP1 targets contributes to the loss of self-renewal. This therapeutic activity extended to primary *SETBP1*-mutant leukemia patient samples, where ziftomenib, revumenib and bleximenib impaired clonogenic growth, promoted myeloid differentiation and suppressed canonical *SETBP1* target genes (**Figure 5**). Although higher inhibitor concentrations were required than those typically effective in KMT2A-rearranged leukemia cell models, MENIN inhibition exerted minimal effects on normal HSPCs, suggesting a favorable therapeutic window. Collectively, these findings indicate that MENIN inhibition in *SETBP1*-mutant leukemia primarily disrupts stemness and self-renewal and promoting differentiation rather than acute cytotoxicity. This may explain why a prior study^17^ did not identify MENIN inhibition as a dependency in their *SETBP1-*mutant models.

Several important considerations remain. The limitations of this study include the following: although supported across primary human HSPCs, multiple experimental systems, and patient samples, our findings require validation in larger patient cohorts and *in vivo* models. *SETBP1* mutations frequently co-occur with alterations in signaling pathways, epigenetic regulators, and splicing factors that may influence sensitivity to MENIN inhibition^1–3^. In addition, although MENIN inhibition effectively disrupted leukemogenic programs in our models, acquired resistance has been reported in other leukemia subtypes^5,26,45,46^, suggesting that combination approaches may be required to achieve durable responses. Given our finding that SETBP1 physically and functionally interacts with components of the

KMT2A regulatory machinery, targeting this complex from multiple angles may represent an attractive therapeutic strategy. Indeed, WDR5 inhibition has shown efficacy in cells with *SETBP1* overexpression^17^. Although our findings establish a functional interaction between mutant *SETBP1* and the KMT2A-MENIN complex, the mechanism by which *SETBP1* mutations enhance KMT2A occupancy remains unclear. Recent studies in *NPM1*-mutant leukemia suggest that MENIN-associated transcriptional machinery can assemble into higher-order condensates that concentrate leukemogenic regulators at target loci, raising the possibility that mutant *SETBP1* contributes to the formation or stabilization of similar regulatory hubs^44–46^. Defining how mutant *SETBP1* promotes KMT2A-MENIN activity and how resistance emerges following MENIN inhibition may also inform rational combination strategies targeting complementary *SETBP1* dependencies, including XPO1, a critical vulnerability in SETBP1-mutant leukemia^47^.

In summary, we identify a previously unrecognized dependence on the KMT2A-MENIN complex in *SETBP1*-mutant leukemias. Mutant *SETBP1* establishes a permissive chromatin landscape that sustains *HOXA/MEIS1*-driven leukemogenic programs and self-renewal. Genetic and pharmacologic disruption of the KMT2A-MENIN complex dismantles these programs, induces differentiation, and impairs clonogenic growth in both experimental cell models and patient leukemias. Together, these findings provide mechanistic insight into *SETBP1*-driven leukemogenesis and support the clinical evaluation of KMT2A-MENIN interaction inhibitors, alone or in rational combination strategies, for patients with *SETBP1*-mutant myeloid malignancies.

## ACKNOWLEDGEMENTS

Funding for this work was provided by a Blood Cancer United Scholar Award, a Vera and Joseph Dresner Foundation MDS Research Fund Established Investigator Award, NIH/NHLBI R01 HL157147, the Andrew McDonough B^+^ Foundation Childhood Cancer Research Grant, a Kuni Foundation Imagination Award, the Knight Cancer Institute NCI Cancer Center Support Grant (P30CA069533), an Edward P. Evans Foundation Discovery Grant, a V Foundation for Cancer Research Grant and the Alex’s Lemonade Stand Foundation Grant (1533610) to JEM. ST was supported by a Blood Cancer United Fellow Scholar Award and Medical Research Foundation Early Clinical Investigator grant. The authors thank the following Oregon Health and Science University core facilities for their assistance: OHSU Flow Cytometry and Monoclonal Antibody Shared Resource (RRID:SCR_009974) and sequencing services were performed by the OHSU Massively Parallel Sequencing Shared Resource. The research reported in this publication used computational infrastructure supported by the Office of Research Infrastructure Programs, Office of the Director, of the National Institutes of Health under Award Number S10OD034224. The content is solely the responsibility of the authors and does not necessarily represent the official views of the National Institutes of Health. We thank Kura Oncology for generously providing ziftomenib/KO539 for use in these experiments. We thank Peter van Galen for providing us with the lentiviral pSMAL-Luc construct. The authors sincerely thank the patients who participated in the BeatAML studies, as well as the clinical and laboratory research teams at the participating academic institutions for their contributions to sample collection and processing.

## AUTHORSHIP CONTRIBUTIONS

ST and JEM were responsible for project conception and design. Data collection and analysis were performed by ST, TTN, HLC, and AP. TPB provided supervision and assistance in planning studies. ST and JEM wrote the manuscript. All authors reviewed and revised the manuscript.

