## Supplemental Data for "The KMT2A-MENIN complex is a therapeutic vulnerability in SETBP1-mutant leukemia"

#### METHODS

##### Culture of CD34⁺ primary human bone marrow stem and progenitor cells

Healthy human de-identified bone marrow cells were purchased from AllCells or Stem Cell Technologies. $CD34^+$ cells were cultured in expansion media StemSpan SFEM II (StemCell Technologies, Cat. #09605) supplemented with 10% StemSpan $CD34^+$ Expansion Supplement (StemCell Technologies, Cat. #02691) and 0.2% UM729 (StemCell Technologies, Cat. #72332).

##### Culture of MOLM-20 cells

MOLM-20 cells were obtained from DSMZ ACC 591 and were routinely tested for mycoplasma contamination. Cells were cultured in 80% RPMI (Gibco, #1187509), 20% FBS (Cytiva HyClone, #SH3039603HI) and 1% Pen/Strep (Gibco, #15140122). Cell line identity was not independently authenticated during this study.

##### Production of lentiviral particles in HEK293T-LX cells

HEK293T-LX cells (Clontech/Takara, #632180) expressing the SV40 T-antigen for high transfectability were seeded at an optimal density of 2-4x$10^6$ cells on a 10 $cm^2$ tissue culture plate in DMEM (Gibco, #11965-092) supplemented with 10% FBS (Cytiva HyClone, #SH3039603HI) and 1% Pen/Strep (Gibco, #15140122) to obtain logarithmic cell growth at the time of transfection with approximately 80% confluence. Packaging plasmids psPAX2 (Addgene, #12260) and VSV-G (Addgene, #8454) were added at a 2: ratio (3.4 ug psPax2 and 1.6 ug VSV-G). Transfer plasmids were used at 5 ug per dish. Transfer plasmids included codon-optimized human $SETBP1^{WT}$ or $SETBP1^{D868N}$, which were subcloned into a constitutive lentiviral vector: pSMAL (Addgene, #161785).

Per transfection reaction 60µl of FuGENE® 6 transfection reagent (Promega, #E269) and 540µl of Opti-MEM (Gibco #31985062) were incubated with the plasmid mixture for 15 minutes at room temperature before adding to the cells to facilitate delivery into cells. Transfected HEK 293T-LX cells were incubated at 37°C for 48h. Viral supernatant was harvested 48h after transfection, passed through 0.45 µm nylon syringe filters (Thermo Scientific, #727-2045), and concentrated in Vivaspin-20 centrifugal concentrators (Sartorius, Cat. VS2042, LOT: 211800217) at 3000rpm at 4°C until a final volume of 500 ul, corresponding to a 40x fold concentration. Viral supernatant was used freshly.

##### Viral transduction of human CD34⁺ primary cells

0.4-0.7 x $10^{\wedge 6}$ primary $CD34^+$ bone marrow cells were spin-infected in expansion medium with 40-fold concentrated lentivirus in the presence of 5µg/ml polybrene for 90 minutes at 1000 g and 32°C. 24 hours after transduction, cells were washed twice with PBS and resuspended in fresh expansion medium. 48-72 hours post-transduction, positive GFP cells were sorted on a SH800S Cell Sorter (Sony Biotechnology) and plated into fresh expansion medium.

##### Colony-forming unit assays

10,000 $CD34^+$ *SETBP1*-mutant positive GFP cells were sorted directly into 4ml MethoCult H4435 (StemCell Technologies, Cat. #04435) supplemented with 1% Penicillin-Streptomycin and 0.1% Fungizone. Primary patient samples were thawed and immediately plated at 2000-5000 cells per mL for $CD34^+$ enriched cells or 100,000-150,000 cells per mL for whole bone marrow. Cells were mixed by vigorous vortexing, and 1 mL of media and cells were plated in triplicate in a 6-well SmartDish (StemCell Technologies, Cat. #27370). After incubating for 14 days, plates were imaged on a STEMvision (StemCell Technologies). For replating of $CD34^+$ *SETBP1*-mutant HSPCs, cells were washed with IMDM supplemented with 20% FBS before plating 25,000 cells per well. Colonies were counted automatically and manually in ImageJ after being blinded by another investigator.

**CRISPR-Cas9 editing**

$CD34^+$ cells were sorted and grown in expansion medium for 7 days after transduction. CRISPR-Cas9 mediated knockout was performed using a Lonza P3 Primary Cell 4-D Nucleofector X Kit S (Cat. # V4XP-3032) in a Lonza 4D-Nucleofector X Unit (Cat. # AAF-1003X).

RNP complexes were prepared by diluting triplet sgRNAs (EditCo Bio, Inc.) to a working concentration of 30 µM. sgAAVS1 #4 XDel (EditCo Bio, Inc.) was used for safe harbor locus targeting. For each nucleofection reaction, 180 pmol of sgRNA and 20 pmol of SpCas9 were combined in P3 buffer supplemented according to the manufacturer's instructions. RNA complexes were incubated for 15 minutes at room temperature to allow for complex assembly.

Following incubation, 125,000 $CD34^+$ cells per reaction were resuspended in nucleofection solution containing RNP complexes and transferred to nucleocuvette wells. Cells were nucleofected using the DZ-100 program. A recovery plate with pre-warmed expansion medium was prepared in advance. Immediately after nucleofection, cells were gently resuspended in medium and transferred from nucleocuvette wells to the recovery plate. Cells were returned to standard culture conditions, maintained in expansion medium and counted every other day.

| Gene | Triplet sgRNA guides |
|---|---|
| MEN1 | GCCCGCCGCCAUGGGGCUGA,<br>UGCUGCCGAGCUGGGCCGAG,<br>GGGCUUCGUGGAGCAUUUUC |
| KMT2A | UGACGAAGACGAAGACGAGG,<br>CGUCGAAGCCCGGGCCCACC,<br>CGCUCUCCCCAAACACGGCC |
| KMT2B | CAACCGCCUCACCUCCUCGU,<br>GCCGCCUGUGGGCCGGCCCG,<br>CCCGAGUCGAGGCUGCGUGC |
| DOT1L | GGAGACCUCCUUCGACCUGG,<br>CGCUCCCCAAGUCCACAAAC,<br>UCUCGGGGUCGGUCACCGAG |

**RNA isolation, cDNA synthesis, and quantitative RT-PCR analysis**

Briefly, cells from liquid culture or CFU assays were isolated and washed well with PBS. Total cellular RNA was isolated with Quick-RNA Microprep Kit (ZymoResarch, #R1051) according to the manufacturer's instructions. 500 ng of total RNA was used for first-strand cDNA synthesis using a High-Capacity cDNA reverse transcription kit (ThermoFisher, #4374966) following the manufacturer's protocol. The program was run for 10 min at 25°C, 2 hours at 37°C, 5 seconds at 85°C, and infinite 4°C. The resulting cDNA was diluted 1:5 and quantitative RT-qPCR was performed on QuantStudio 7 Flex Real-Time PCR System (Applied Biosystems) using Taqman probes (ThermoFisher) for MECOM (Hs00602795), HOXA9 (Hs04931836), HOXA10 (Hs00172012), MEIS1 (Hs00180020), MYC (Hs00905030) and GUSB (Hs00939627) with TaqMan™ Universal PCR Master Mix (ThermoFisher, 4304437). Ct values were normalized to *GUSB* expression and relative expression was quantified using the $2^{-\Delta\Delta CT}$ method.

**RNA sequencing and analysis**

RNA was extracted as described above. mRNA libraries were prepared with the Takara Smart-Seq V4 Plus kit (TakaraBio, #R400752) according to the manufacturer's protocol. 10 ng of RNA for each sample was used for mRNA capture. Libraries were sequenced on the Illumina Nova Seq 6000 with 2x 150 bp paired-end reads.

Reads were trimmed using Fastp[48] with options '--detect_adapter_for_pe', '--trim_poly_g', and '--adapter_fasta' using a custom FASTA file of adapter sequences provided by the library kit manufacturer. Trimmed reads were aligned to the human genome (GRCh38/hg38) using STAR[49]. A raw gene-level counts matrix was generated from STAR-aligned reads, and DESeq2[27] was used for normalization and differential expression analysis. Transcription factor activity was predicted from normalized RNA-seq counts using Priori[32]. Gene Set Enrichment Analysis (GSEA)[28,29], was performed on a pre-ranked gene list ordered by the Wald statistic outputted by DESeq2. GSEA was run with the CGP (chemical and genetic perturbation) subset from the C2 curated gene sets and Hallmark gene sets from the Molecular Signatures Database (MSigDB)[50].

**CUT&Tag**

Cells were counted, and 100,000 cells per antibody per condition were harvested by centrifugation at 300g. Cells were then washed in 1 mL PBS before being resuspended in 1 mL cold NE2B buffer (25 mM HEPES-KOH pH 7.9, 12.5 mM KCl, 0.125% Triton X-100, and 0.5 mM spermidine trihydrochloride (MP Biomedicals, Cat. #0210047201) in molecular grade water) and incubated for 10 minutes to extract nuclei on ice. Nuclei were pelleted and resuspended at $1 \times 10^6$ nuclei per mL in cold NE2B buffer. For each sample, 100 µL of nuclei were mixed with 10 µL of ConA beads (Bangs Laboratories, #NC1781147) pre-washed three times in ConA-Binding buffer (20 mM HEPES-KOH pH 7.9, 10 mM KCl, 1 mM CaCl2, and 1 mM MnCl2 in molecular grade water) and incubated for 10 minutes. Bead-bound nuclei were then immobilized on a magnet and incubated with 1:100 diluted primary antibodies (KMT2A/MLL1C (Cell Signaling Technology, Cat. #14197), MENIN (Cell Signaling Technology, Cat. #19893), H3K4me3 (Cell Signaling Technology, Cat. #9751S), H3K27ac (Abcam, Ab4729), H3K27me3 (Cell Signaling Technology, Cat. #9733), SETBP1 (ProteinTech, #16841-1-AP), Rabbit IgG (Cell Signaling Technology, Cat. #2729) in antibody buffer (20 mM HEPES pH 7.5, 150 mM NaCl, 0.5 mM spermidine trihydrochloride, 2 mM EDTA, 1X protease inhibitor (Pierce mini-tablets EDTA-free, Cat. #A32955), and 0.01% digitonin in molecular grade water) on a nutator overnight at 4 °C.

The next day, samples were magnetically separated and incubated with secondary antibodies (Guinea pig anti-rabbit (Antibodies Online, Cat. #ABIN101961) diluted 1:100 in cold Digitonin-150 buffer (20 mM HEPES pH 7.5, 150 mM NaCl, 0.5 mM spermidine trihydrochloride, 1X protease inhibitor, and 0.01% digitonin in molecular grade water). Samples were then placed on a nutator at room temperature for 30 minutes. After incubation, samples were washed twice with 200 µL of cold Digitonin-150 buffer before resuspending in 50 µL of Digitonin-300 buffer (20 mM HEPES pH 7.5, 200 mM NaCl, 0.5 mM spermidine trihydrochloride, 1x protease inhibitor, and 0.01% digitonin in molecular grade water) with 2.5 µL of pA/G-Tn5 (Epicypher Cat. #15-1017) and gently vortexing to mix. Samples were then placed on a nutator at room temperature for 1 hour. After incubation, samples were washed twice with 200 µL cold Digitonin-300 buffer and the supernatant was removed. Samples were resuspended in 50 µL cold Tagmentation buffer (Digitonin-300 buffer supplemented with 10 mM MgCl2) and incubated in a thermocycler for 1 hour at 37 °C (lid temperature 47 °C). Samples were then placed on a magnet stand, and the supernatant was removed before washing with 50 µL of TAPS Wash buffer (10 mM TAPS pH 8.5 and 0.2 mM EDTA in molecular grade water). DNA was released using 5 µL SDS release buffer (0.1% SDS and 10 mM TAPS pH 8.5 in molecular grade water) and vortexing before incubating in a thermocycler at 58 °C for 1 hour (lid temperature 68 °C). Samples were removed from the thermocycler and 15 µL of Triton neutralization buffer (0.67% Triton-X100 in molecular grade water) was added. Samples were PCR amplified using custom primers at 1 uM and NEBNext High Fidelity 2X PCR mix (New England Biolabs, Cat. #M0541) under the following conditions: 5 minutes at 58 °C, 5 minutes at 72 °C, 45 seconds at 98 °C, 16 cycles of 15 seconds at 98 °C and 10 seconds at 60 °C, and 60 seconds at 72 °C. Samples were purified using Omega Bio-Tek Mag-Bind TotalPure NGS beads at 1.3x volume before elution in 10 mM Tris-HCl pH 8.0. Samples were quantified using a Qubit fluorometer (1X dsDNA High Sensitivity Kit (Invitrogen, Cat. #Q33230)) and on a TapeStation (HS D1000 Kit) before pooling and sequencing on an Illumina NextSeq 2000 with 50 or 75 bp paired-end reads.

**CUT&Tag Analysis**

Reads were trimmed using Fastp[48] with options "--detect_adapter_for_pe", "--trim_poly_g", and "--adapter_fasta" using a custom FASTA file of adapter sequences provided by the library kit manufacturer. Trimmed reads were aligned to the human genome (GRCh38/hg38) using Bowtie2[51] with parameters "--local --very-sensitive-local --no-unal --no-mixed --no-discordant --phred33 -I 10 -X 700". Duplicates were marked using the Sambamba[52] "markdup" subcommand. Peaks were called using GoPeaks[53] using IgG samples as negative controls. To define consensus regions for each antibody target, peaks present in at least two replicates per sample condition were retained as high-confidence regions. These reproducible peaks were then merged across conditions using BEDTools[54] to generate a unified consensus peak set for each target. High-confidence regions co-occupied by multiple antibody targets were also determined using BEDTools. ChIPseeker[55] and TxDb.Hsapiens.UCSC.hg38.knownGene were used to annotate binding regions to genomic features and determine the percentage of regions in each feature.

Signal at consensus regions was quantified and analyzed using BEDTools and deepTools[56]. Counts-per-million (CPM) normalized bigWig tracks were generated using deepTools bamCoverage with parameters "--binSize 10 --smoothLength 50 --normalizeUsing CPM". Coverage tracks were then averaged across replicates within each sample group using WiggleTools[57] and the UCSC bedGraphToBigWig tool[58]. To generate heatmaps and median signal profiles of normalized CUT&Tag signal in regions spanning +/- 3kb from consensus region centers, we used deepTools computeMatrix with parameters "--referencePoint center --upstream 3000 --downstream 3000" and deepTools plotHeatmap with parameter "--averageTypeSummaryPlot median".

To predict SETBP1 target genes, a Python 3 implementation (https://github.com/crazyhottommy/BETA2) of the BETA[31] algorithm (with default thresholds) was used on all SETBP1 consensus peaks and differential gene expression data from *SETBP1*$^{D868N}$ and *Luc*-control HSPCs. For each gene, a regulatory potential score was calculated based on the distance-weighted contribution of nearby peaks, and a rank product integrating both regulatory potential and differential expression was generated. Genes with a rank product <0.001 were considered high-confidence targets. Nominated target genes were compared to leading edge genes from GSEA gene sets that were significantly enriched in *SETBP1*$^{D868N}$ cells (positive normalized enrichment score, FDR <0.05). Overlap between targets and leading edge genes was assessed with a one-sided hypergeometric test using the GSEA ranked gene list as background, and Benjamini-Hochberg (BH)[59] correction was applied to p-values.

**Flow cytometry**

0.2-0.5 x $10^6$ cells were washed, resuspended in BD Pharmingen™ Stain Buffer (FBS) (BDBiosciences, Cat #554656) and incubated with fluorochrome-conjugated antibodies (CD11b, CD15, CD14, CD117, HLA-DR) for 30 min at 4°C. Following this, cells were washed with PBS to remove nonspecific antibody binding. Stained cells were resuspended in Stain buffer, and data was collected from each cell and assessed in two-dimensional plots on LSR Fortessa (BD, New Jersey, USA). Further analysis was performed with FlowJo software (Tree Star). Population gating was always performed according to an unstained sample.

**Menin inhibitor studies**

Ziftomenib was obtained from Kura Oncology. Revumenib (SNDX-5613 Cat. No. S8996) and Bleximenib (Cat. No. E4679) were purchased from Selleck Chemicals. Cells were treated with the indicated concentrations of MENIN inhibitor or control (0.02% DMSO). For CFU assays, treated cells were plated in MethoCult H4435 (StemCell Technologies, Cat. #04435) supplemented with 1% Penicillin-Streptomycin and 0.1% Fungizone. Colonies were counted with StemVision after 14 days. Where indicated, cells were serially replated to assess self-renewal capacity. *SETBP1*-mutant primary leukemia patient samples could not be replated. Colonies were counted either using the STEMvision

Analyzer software, or images were blinded and manually counted in ImageJ after being blinded by another investigator.

For liquid-culture experiments, cells were culture in expansion medium: StemSpan SFEM II (StemCell Technologies, Cat. #09605) supplemented with 10% StemSpan CD34+ Expansion Supplement (StemCell Technologies, Cat. #02691) and 0.2% UM729 (StemCell Technologies, Cat. #72332). Cells were collected by centrifugation every 48 hours and resuspended in fresh expansion medium containing freshly prepared MENIN inhibitor or corresponding DMSO control. Cells were harvested at the indicated experimental time points to assess gene expression, immunophenotype, RNA sequencing or CUT&Tag analysis.

##### *SETBP1*-mutant leukemia patient samples

Frozen peripheral blood or bone marrow cells were obtained from six patients diagnosed with either CNL, aCML, or CMML-AML after informed consent on IRB protocol number 4422. Frozen vials were rapidly thawed in a 37°C water bath. Cells were then treated with DNase I (STemCell Technologies, Catalog #07900) following the manufacturer's instructions to prevent clumping caused by release of genomic DNA. Detailed clinical and molecular characteristics, including co-occurring mutations and cell density for CFU assays, are described below. After scoring colonies, cells were harvested from patient 23-00049, 24-00602 and 17-00285 for flow cytometric immunophenotyping. For gene expression experiments cells from colonies were harvested from patient 16-00130, 16-00254, 23-00049, 17-00285.

| Patient ID | Source | Type | Diagnosis | Mutation | Alteration | Co-occurring mutations | Cells plated in Methocult total (4ml) | Cells per ml |
|---|---|---|---|---|---|---|---|---|
| 24-00602 | Bone Marrow | CD34+ selected | CNL | SETBP1 | G870S | CSF3R_T618I, SRSF2_P95_102del, ASXL1_W796*, ETV6_R163Qfs*6, PTPN11_N308D, JAK1_S94A, NPM1_R197Q, BRAF_L745V | 20,000 | 5,000/ml |
| 22-00633 | Bone Marrow | CD34+ selected | CMML - AML | SETBP1 | R942W | SRSF2_P95H, ASXL1_G646fs*12, CBL_C384Y, RUNX1_A156E, TET2_G1861V, GATA2_V211L, DNAH5_W3350L | 10,000 | 2500/ml |
| 23-00049 | Peripheral Blood | CD34+ selected | aCML | SETBP1 | D868N | CUX1_Y935*, SRSF2_P95H, PTPN11_N308D, ASXL1_G646Wfs*12, TET2_R550, TET2_R1440Tfs*38, NF1_H130R | 8,000 | 2,000/ml |
| 16-00254 | Bone Marrow | CD34+ selected | CNL | SETBP1 | G870S | CSF3R_T618I, SRSF2_P95R, ASXL1_G646fs*12, MPL_R102P, STAG2_splice | 17,000 | 4250/ml |
| 17-00285 | Peripheral Blood | not selected | CNL | SETBP1 | G870S | ASXL1_G646fs*12, CSF3R_Q776, CSF3R_T640N, TCF3_R574H | 600,000 | 150,000/ml |
| 16-00130 | Bone Marrow | not selected | aCML | SETBP1 | D868N | SRSF2_P95L, NRAS_G12R, GATA2_M325fs*2, ASXL1_E635fs*15 | 400,000 | 100,000/ml |

### Suppl. Fig. 1

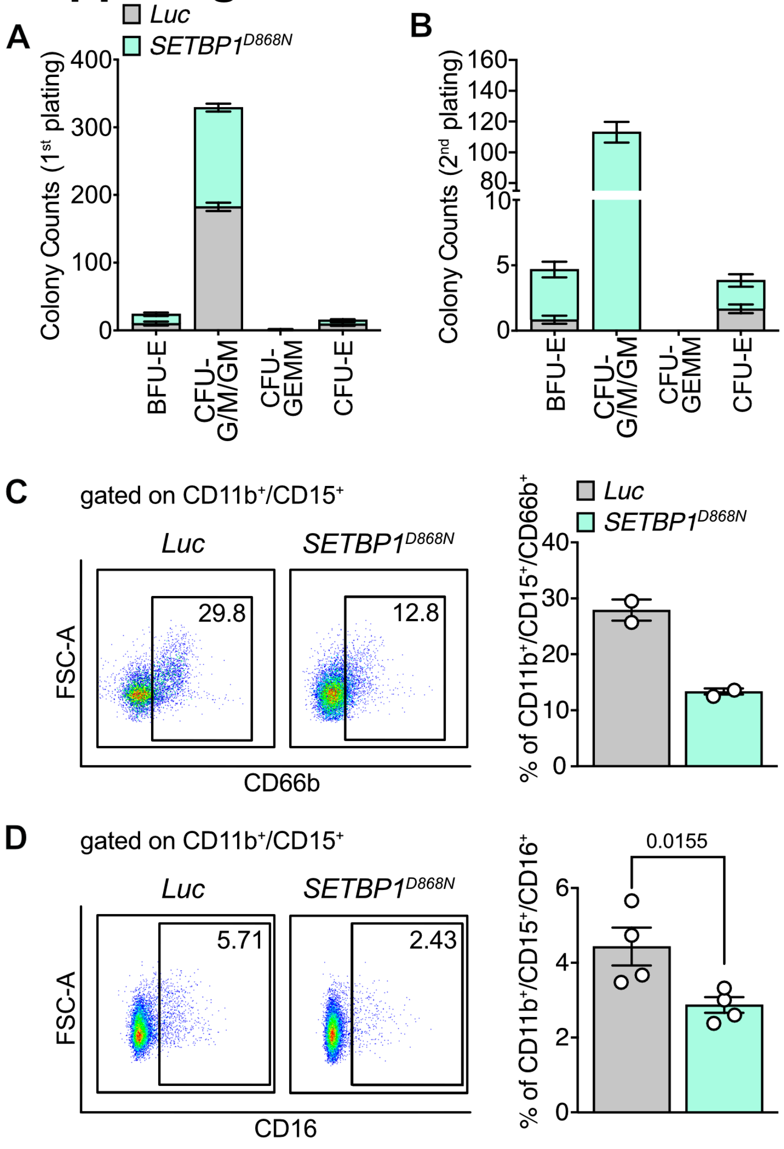


**Supplemental Figure 1. A-B.** Quantification of automated colony-forming unit counting and characterization of colony morphology of Luc and *SETBP1*$^{D868N}$ expressing HSPCs. CFU-GEMM, CFU-GM, and BFU-E cells were counted after 1$^{st}$ (A) and 2$^{nd}$ (B) plating. **C-D.** Representative flow cytometry images and quantification of *Luc* and *SETBP1*$^{D868N}$-expressing cells cultured for 13 days after sorting are shown. CD66b$^{+}$ (C) and CD16 (D) cells were gated on CD11b$^{+}$/CD15$^{+}$ cells.

### Suppl. Figure 2

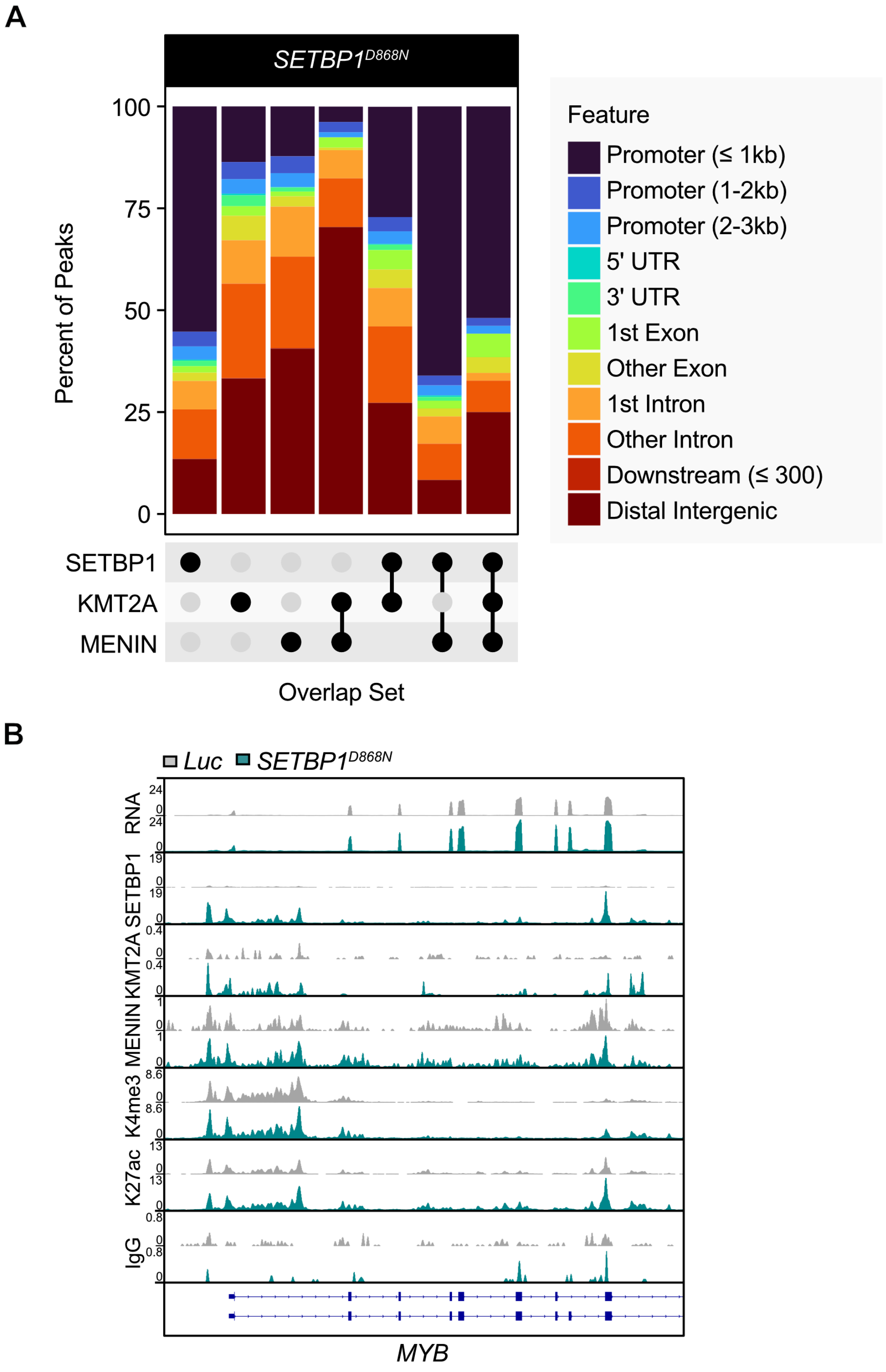


**Supplemental Figure 2. A.** Genomic feature distribution for unique peaks in SETBP1, KMT2A, MENIN, and shared (co-occupied) regions in *SETBP1*$^{D868N}$ HSPCs. Approximately 55% of the shared regions overlap with promoter regions. **B.** Merged IGV tracks for RNA-seq expression data (n=5 per group), SETBP1 (n=3 per group), KMT2A (n=3 per group), MENIN (n=3 for *SETBP1*$^{D868N}$, n=2 for *Luc*), H3K4me3 (n=3 per group), and H3K27ac (n=3 per group) are shown at the *MYB* locus. IgG serves as a control.

### Suppl. Figure 3

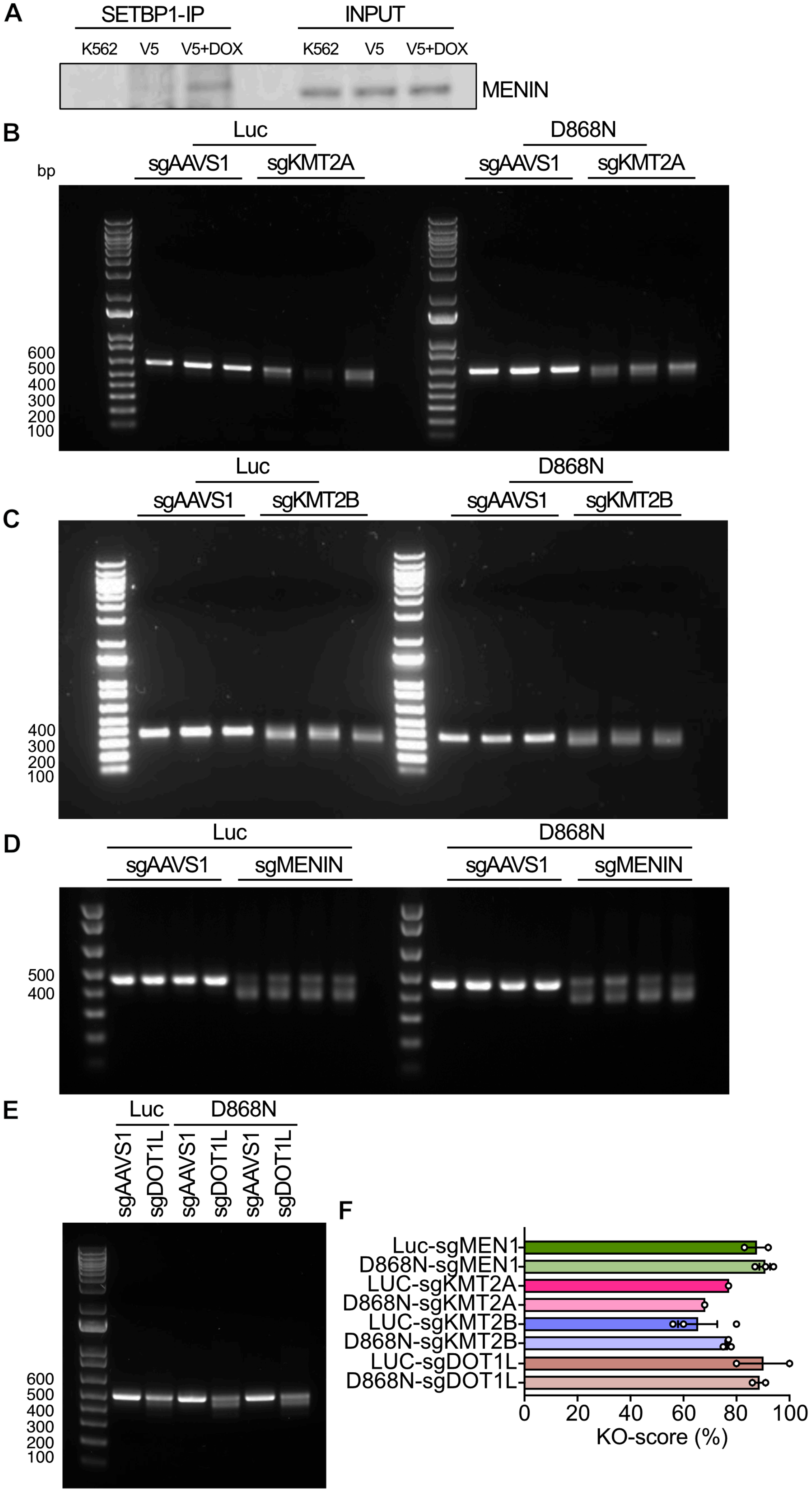

**Supplemental Figure 3. A.** Co-Immunoprecipitation of V5-SETBP1$^{D868N}$ with MENIN in K562 cells. The V5 antibody was used for immunoprecipitation, followed by immunoblotting with anti-MENIN. 5% of the total lysate was run for input. Constructs were induced with doxycycline (+Dox). Untransduced K562 cells (K562) are shown as a control. **B-E.** Validation of CRISPR/Cas9-mediated genome editing 7 days after nucleofection. The Cas9 nuclease was targeted to (B) KMT2A, (C) KMT2B, (D) MENIN, or (D) DOT1L genes by selected triple-sgRNAs in *SETBP1*$^{D868N}$ or *Luc*-control cells. **F.** Inference of CRISPR Editing (ICE) and Knockout (KO) scores for Sanger-sequenced PCR products.

### Suppl. Fig. 4

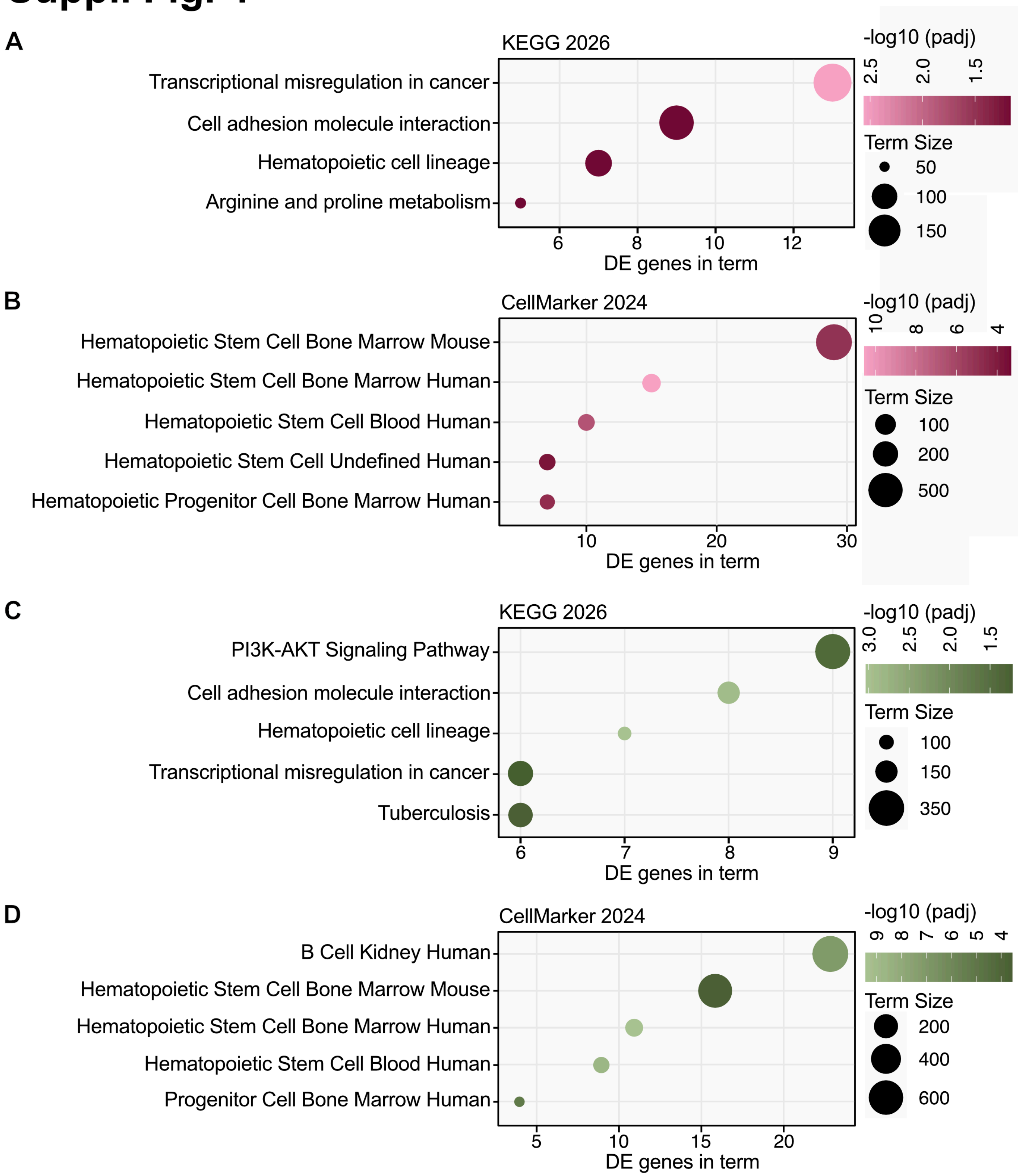


**Supplemental Figure 4. A.** EnrichR analysis of differentially expressed (DE) genes downregulated following KMT2A loss (log2FC < -0.5, padj < 0.05) identified significantly enriched pathways (padj < 0.1) in the KEGG 2026 database. **B.** EnrichR analysis of DE genes downregulated following KMT2A loss (log2FC < -0.5, padj < 0.05). Top 5 most enriched cell type signatures (padj < 0.01) in the

CellMarker 2024 database are shown. **C.** EnrichR analysis of DE genes downregulated following MENIN loss (log2FC < -0.5, padj < 0.05) identified significantly enriched pathways (padj < 0.1) in the KEGG 2026 database. **D.** EnrichR analysis of DE genes downregulated following MENIN loss (log2FC < -0.5, padj < 0.05). Top 5 most enriched cell type signatures (padj < 0.01) in the CellMarker 2024 database.

### Suppl. Fig. 5

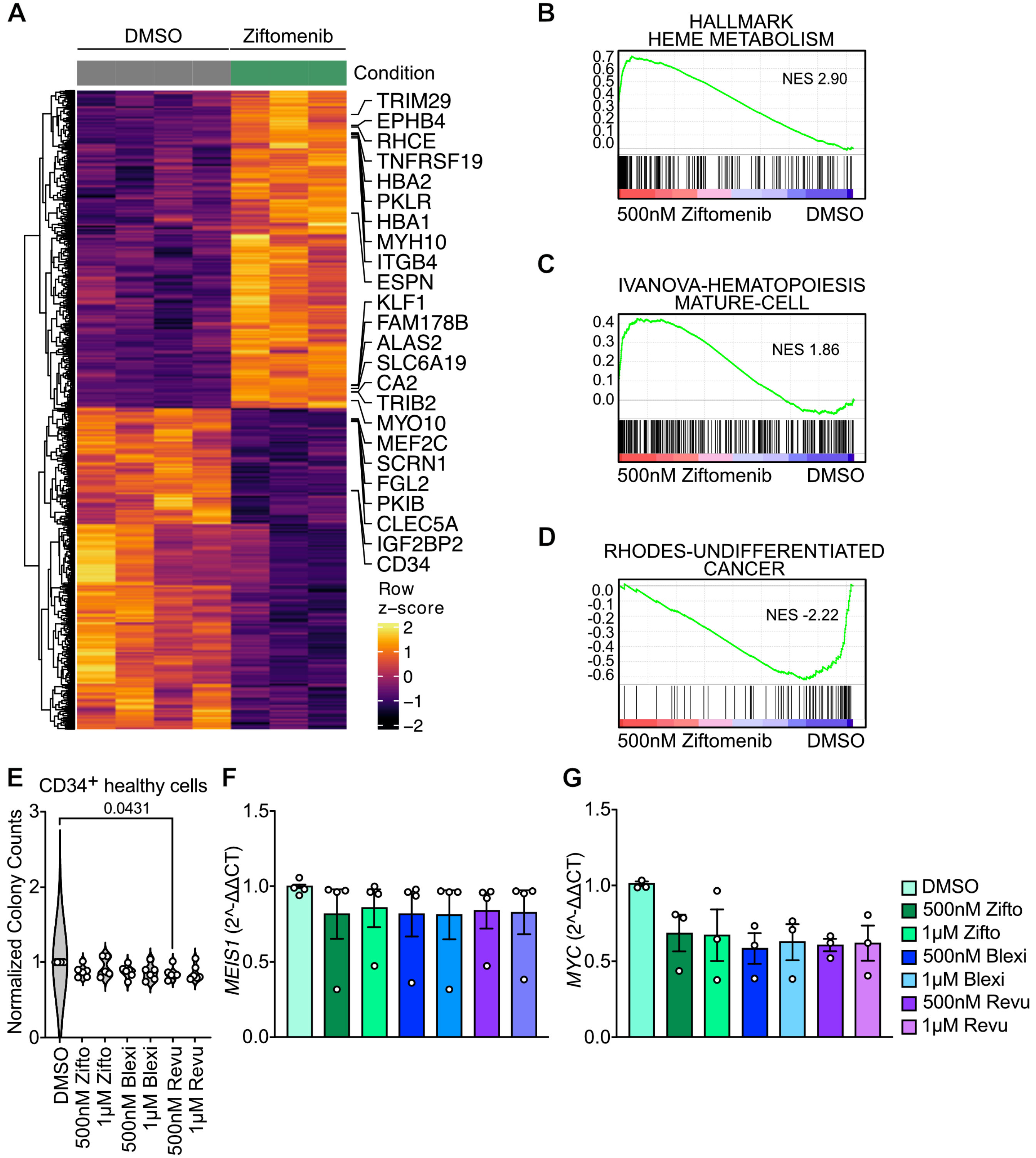


**Supplemental Figure 5. A.** Heatmap of all significantly differentially expressed genes (padj <0.05) in *SETBP1*$^{D868N}$ HSPCs following six days of Ziftomenib treatment. The top 5 most significantly DE upregulated genes and genes among the top 50 that also overlap with the “Genes upregulated during hematopoietic differentiation” gene set (PMID:19128795) from the HDSigDB_Human_2021[60] library are annotated in the Ziftomenib condition. The top 6 most significantly DE upregulated genes and CD34 (ranked #12) in the DMSO condition are annotated. **B-C.** Gene Set Enrichment Analysis (GSEA) of Ziftomenib and DMSO control conditions using C2: CGP and Hallmark gene sets (q-value < 0.05). Positive normalized enrichment scores (NES) for the top gene sets enriched in the Ziftomenib condition. **D.** Gene Set Enrichment Analysis (GSEA) of Zifomenib and DMSO control conditions using C2: CGP and Hallmark gene sets (q-value < 0.05). Negative normalized enrichment scores (NES) for the top gene sets enriched in the DMSO condition. **E.** Normalized colony counts after 14 days of treatment with Ziftomenib, Bleximenib or Revumenib in healthy primary human $CD34^+$ cells. **F-G.** qRT-PCR analysis of *MEIS1 and MYC* mRNA transcripts from colonies harvested at 14 days. Expression levels were normalized to the *GUSB* housekeeping gene and DMSO. Each replicate represents a biologically independent sample from a distinct patient. If not otherwise described, statistical analyses were performed with 2-way ANOVA with Sidak’s multiple comparison test. Error bars represent the standard error of the mean (SEM) for E, F and G.

#### SUPPLEMENTAL TABLES

Table S1: SETBP1-D868N-vs-Luc_RNAseq-DEgenes_padj0.05
Table S2: SETBP1-D868N-vs-Luc_Priori_mean_diff_activity
Table S3: SETBP1-D868N_highConfidence targets_Beta2
Table S4: SETBP1-D868N_beta2_genes_in_gsea_leading_edge_hypergeometric_test
Table S5: D868N_KMT2A-vs-D868N_AAVS1-0.05
Table S6: D868N_MEN1-vs-D868N_AAVS1-0.05
Table S7: D868N_500nM Ziftomenib-vs-DMSO-0.05
